# Nucleic acid sequence contributes to remodeler-mediated targeting of histone H2A.Z

**DOI:** 10.1101/2023.12.06.570360

**Authors:** Cynthia Converso, Leonidas Pierrakeas, Lirong Chan, Shalvi Chowdhury, Vyacheslav I. Kuznetsov, John M. Denu, Ed Luk

## Abstract

The variant histone H2A.Z is inserted into nucleosomes immediately downstream of promoters and is important for transcription. The site-specific deposition of H2A.Z is catalyzed by SWR, a conserved chromatin remodeler with affinity for promoter-proximal nucleosome depleted regions (NDRs) and histone acetylation. By comparing the genomic distribution of H2A.Z in wild-type and SWR-deficient cells, we found that SWR is also responsible for depositing H2A.Z at thousands of non-canonical sites not directly linked to NDRs or histone acetylation. To understand the targeting mechanism of H2A.Z, we presented SWR with a library of nucleosomes isolated from yeast and characterized those preferred by SWR. We found that SWR prefers nucleosomes associated with intergenic over coding regions, especially when polyadenine tracks are present. Insertion of polyadenine sequences into recombinant nucleosomes near the H2A-H2B binding site stimulated the H2A.Z insertion activity of SWR. Therefore, the genome is encoded with information contributing to remodeler-mediated targeting of H2A.Z.

## INTRODUCTION

Nucleosomes are the building blocks of chromosomes in a eukaryotic cell [1]. A typical nucleosome has a protein core made up of eight histones, two copies of histone H2A, H2B, H3 and H4, that is coiled around by ∼146-basepair (bp) DNA in 1.65 superhelical turns [2]. Individual nucleosomes are connected by linker DNA of variable lengths and are positioned along genes in a non-random, stereotypic pattern that is functionally important for transcription [3]. In yeast, a nucleosome-depleted region (NDR), 80-200 bp in length, is typically associated with a promoter [4]. Flanking the NDR are the +1 and -1 nucleosomes. In yeast, the upstream edge of a +1 nucleosome overlaps the transcription start site (TSS) [5]. Downstream of the +1 nucleosome are the +2, +3 nucleosomes and so on, organized into a closely spaced array [4]. Since the nucleosome structure generally occludes DNA elements, focused assembly of the transcriptional preinitiation complex (PIC) occurs at the NDR [6]. Studies in yeast showed perturbations of the position and occupancy of the native chromatin arrangement led to aberrant transcriptional response and initiation from cryptic promoters within genic regions, negatively impacting the fitness of cells [7–10].

The +1 nucleosome is frequently installed with the histone variant H2A.Z, a prominent landmark associated with active and poised promoters [5,11]. H2A.Z is required for rapid transcriptional response in yeast and is essential for life in metazoans [12–15]. RNA polymerase II (Pol II)-mediated transcription preferentially ejects H2A.Z over H2A, suggesting that an H2A.Z-containing nucleosome at +1 is the chromatin state designated for initiation [16,17]. However, the role of H2A.Z appears to extend beyond transcription initiation. In yeast, optimal phosphorylation of Pol II C-terminal domain (CTD), elongation factor recruitment, Pol II elongation rate, and RNA splicing require H2A.Z [18,19]. In metazoans, H2A.Z nucleosomes at +1 sites regulate the pause release of Pol II [20,21]. These studies suggest that H2A.Z functions at a juncture whereby Pol II transitions from initiation to productive elongation.

The site-specific deposition of H2A.Z is carried out by SWR, a 14-component chromatin remodeling complex [22–24]. The ‘SWR’ nomenclature distinguishes the complex from its core subunit, Swr1, which is a member of the SWI/SNF-related family ATPases [25]. Unlike other chromatin remodelers in the SWI/SNF family, which slide nucleosomes, SWR catalyzes an ATP-driven remodeling reaction that installs H2A.Z into nucleosomes. At the molecular level, SWR replaces the two H2A-H2B (A-B) dimers within a canonical H2A-containing (AA) nucleosome with free H2A.Z-H2B (Z-B) dimers that are delivered by histone chaperones in a stepwise, distributive manner [26,27]. As such, SWR produces a heterotypic H2A/H2A.Z (AZ) nucleosome as an intermediate before forming the homotypic H2A.Z (ZZ) nucleosome as the final product [28]. The targeting of SWR is mediated in part by the Swc2 subunit, which is a DNA binding protein [29]. Swc2 guides SWR to the NDR by facilitating a one-dimensional search for long linker DNA along chromatin [30]. Histone acetylation is also implicated in SWR recruitment and H2A.Z deposition [11,31]. SWR bears a tandem bromodomain in the Bdf1 subunit and a YEATS domain in the Yaf9 subunit that are reader modules for acetylated histone tails [11,32,33]. However, SWR’s affinity for the NDR and histone acetylation cannot fully explain the genomic distribution of H2A.Z in vivo, as many H2A.Z-containing nucleosomes are located at NDR-distal sites and are not particularly enriched for histone acetylation based on published datasets [5,34]. Therefore, the targeting of H2A.Z and the contribution by SWR-independent pathways remain incompletely understood.

The H2A.Z insertion activity of SWR is influenced by the DNA sequences of the nucleosomal substrates. When presented with nucleosomal substrates assembled with the Widom ‘601’ DNA sequence, which is asymmetric on either side of the dyad axis, SWR inserts H2A.Z onto the opposite faces of the nucleosome at differential rates [28]. The determinant of the insertion bias was pared down to a 16-bp region in front of the remodeling ATPase engagement site, where the favored sequence has longer poly(dG:dC) tracks (but equal GC content) [28]. However, the sequence variation between the two halves of the Widom sequence is too limiting to truly understand the potential effects that DNA sequence has on the remodeling activity of SWR.

An emerging theme in the understanding chromatin remodeling mechanism is that the effect of DNA sequences on remodeling activity is biologically relevant. How DNA sequences affect the activity of RSC, another chromatin remodeler of the SWI/SNF-related family, is a case in point. RSC is involved in the widening of NDRs at yeast promoters [35,36]. The nucleosome removal activity of RSC is stimulated by poly(dA:dT) sequences, which are frequently found at promoter-proximal NDRs [35,37]. These findings suggest that the genome is encoded with information that can direct remodeling activity in a site-specific manner, contributing to the organization of native chromatin. Whether the genome is encoded with information that can fine-tune the H2A.Z deposition activity of SWR is unknown.

In this study, we determined the genomic distribution of SWR-dependent H2A.Z by comparing chromatin-bound H2A.Z in WT and *SWC2*-deficient cells. We found that SWR is responsible for depositing H2A.Z not only at NDR-proximal sites, but also at thousands of NDR-distal sites that are not particularly enriched for histone acetylation. This observation motivated us to test the hypothesis that DNA sequences contribute to SWR’s substrate specificity across the yeast genome. We developed an in vitro approach that utilizes mono-nucleosome libraries isolated from yeast cells to interrogate the substrate specificity of SWR. We found that SWR targets a subpopulation of nucleosomes containing sequences that are characteristic of intergenic regions, providing an explanation for how SWR targets at least some of the NDR-distal sites.

## RESULTS

### SWR deposits H2A.Z predominantly at +1 nucleosomes but also at non-canonical sites

Since it was unknown what contribution SWR-independent pathway(s) have on H2A.Z deposition, we first determined the cellular levels of chromatin-bound H2A.Z in wild-type (WT) and *SWC2* deficient (*swc2Δ*) yeast cells. The SWR complex requires Swc2 to deposit H2A.Z [29]; therefore, any H2A.Z nucleosomes observed in *swc2Δ* was assumed to be deposited by SWR-independent mechanisms.

To quantify chromatin-bound H2A.Z, we used an in vivo disulfide crosslinking approach called VivosX [38]. This approach took advantage of yeast mutants bearing cysteine-modified *HTZ1(T46C)* and *HTA1(N39C)* as the sole sources of H2A.Z and H2A, respectively. Given that H2A.Z is the only variant H2A in yeast and that no other cysteines are present in its core histones, a pair of cysteine probes contributed by the cysteine-modified H2A.Z and/or H2A would be found at the interdisk interface of nucleosomes and close enough for disulfide crosslinking (**Figure 1A**) [39]. Formation of irreversible crosslinking adducts were induced by the addition of 4,4′-Dipyridyl disulfide (4-DPS), a cell-permeable, thiol-specific crosslinker [38]. In addition, the *HTZ1(T46C)* gene was fused to a C-terminal 2xFLAG tag to facilitate the resolution of the crosslinking adducts in immunoblotting analysis. As such, the crosslinked adducts of H2A-to-H2A.Z and H2A.Z-to-H2A.Z infer AZ and ZZ nucleosomes, respectively, whereas uncrosslinked H2A.Z infers non-nucleosomal H2A.Z (**Figure 1B**). Band intensities were normalized to histone H4, which serves as a loading control (**Figure 1B-1C**). In WT cells, we estimated that 12% of bulk H2A.Z is in the ZZ configuration, 57% in the AZ configuration, and 31% in non-nucleosomal forms (**Figure 1B-C**). In *swc2*Δ, ZZ nucleosomes decreased to 2.5%, AZ nucleosomes decreased to 34%, while non-nucleosomal H2A.Z increased to 47% (**Figure 1B-C**). Therefore, SWR is responsible for the formation of most ZZ nucleosomes; however, more than half of the AZ nucleosomes are formed by SWR-independent mechanisms.

**Figure 1.**
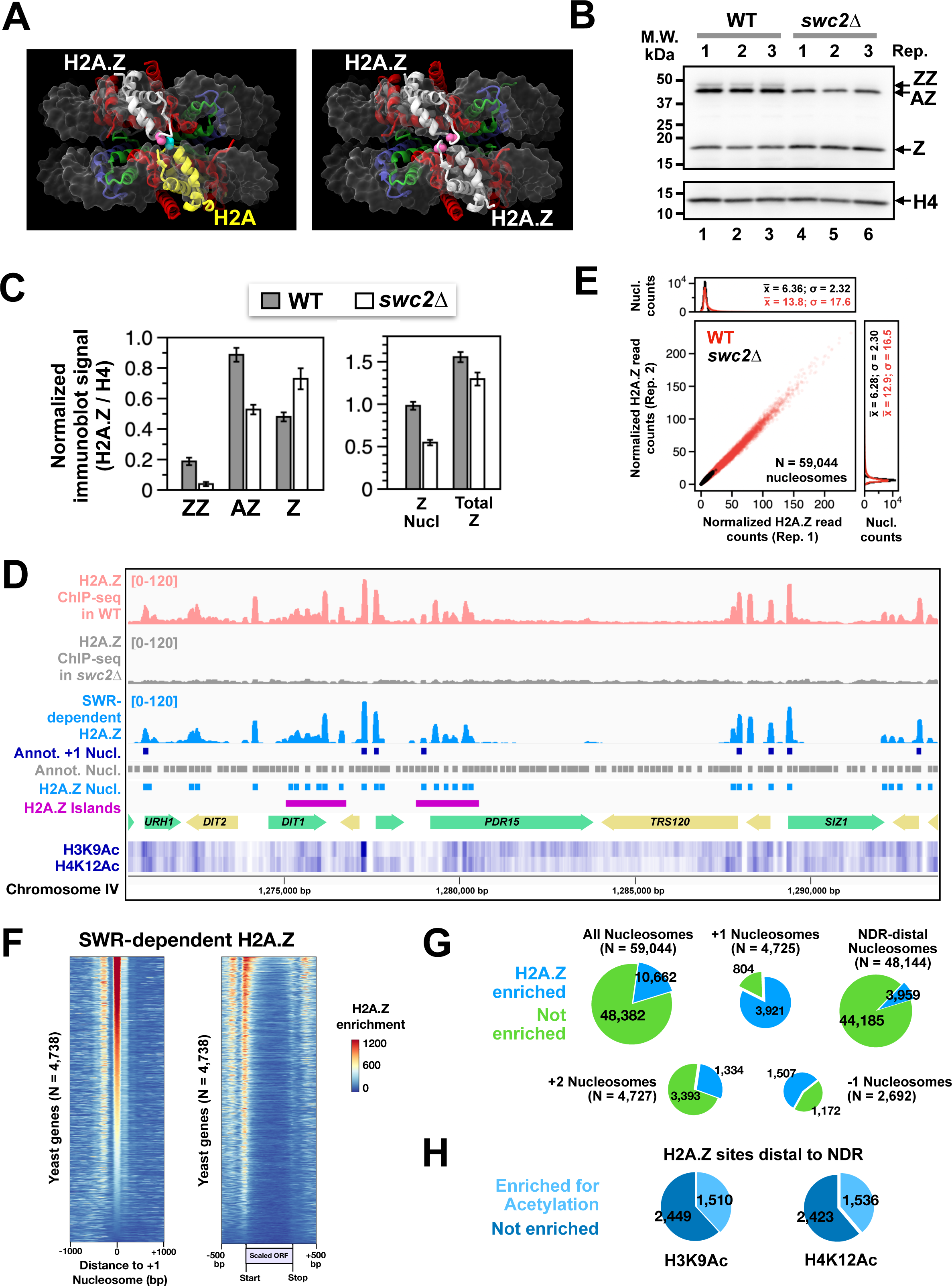
Quantifying SWR-dependent H2A.Z-containing nucleosomes in *S. cerevisiae*. **(A)** The structure of the heterotypic nucleosome containing the human H2A.Z.1 and H2A is on left (PDB: 5B32). The homotypic nucleosome with two copies of H2A.Z.1 is on right (PDB: 5B33). Pink residues in the human H2A.Z.1 that align with T46 of yeast Htz1. Blue residue of human H2A align with N39 of yeast Hta1. **(B)** VivosX analysis of WT and *swc2Δ* cells carrying the *HTZ1(T46C)-2xFLAG* and *HTA1(N39C)* alleles. Total proteins extracted from the indicated strains were analyzed by SDS-PAGE and immunoblotting with anti-FLAG antibody (top) and anti-H4 antibody (bottom). Biological replicates (Rep) represent extracts prepared from individual transformants. **(C)** Quantification of the immunoblots in B. Normalized signals represent band intensities of AZ, ZZ, and Z divided by that of H4. **(D)** ChIP-seq data of H2A.Z in WT (red) and *swc2Δ* cells (black). SWR-dependent H2A.Z signal (blue) is calculated by subtracting *swc2Δ* from WT. Grey boxes: annotated nucleosomes [39,40]. Purple boxes: annotated +1 nucleosomes [40]. Blue boxes: H2A.Z nucleosomes deposited by SWR (this study). Blue heatmaps: H3K9Ac and H4K12Ac [34] **(E)** Read counts of the H2A.Z ChIP-seq data at 59,044 non-repetitive nucleosomal positions were plotted between biological replicates for WT (red) and *swc2*Δ (black) [39,40]. The rectangular boxes above and to the right of the main plot are the histograms of the datapoint densities in the x- and y-axes, respectively. **(F)** Heatmaps of SWR-dependent H2A.Z signals (i.e., WT minus *swc2Δ*) aligned at the center of +1 nucleosomes (left) or at the start and end of open reading frames (ORF) (right). ORFs were scaled using the computeMatrix function in deepTools [48]. **(G)** Pie charts showing the relative number of H2A.Z sites in the indicated categories. **(H)** Pie charts of NDR-distal H2A.Z sites that are enriched for acetylation at H3K9 or H4K12.

To determine the genomic locations of SWR-dependent and -independent H2A.Z nucleosomes, we compared the ChIP-seq profiles of H2A.Z in WT and *swc2Δ* cells. Experimentally, bulk chromatin was isolated from spheroplasted cells followed by fragmentation into mainly mono-nucleosomes by limited micrococcal nuclease (MNase) digestion (**Figure S1A**). The nucleosomes were subjected to immunoprecipitation with anti-FLAG agarose targeting the 2xFLAG epitope tag on the C-terminus of H2A.Z (**Figure S1B**) [16]. The immunoprecipitated DNA fragments were sequenced using the Illumina platform and mapped to the yeast genome (version R64-1-1). Given that H2A.Z-containing nucleosomes (i.e., AZ and ZZ combined) in *swc2*Δ were 55.8% of WT (**Figure 1C**, right), we normalized the H2A.Z ChIP-seq data such that ∼558,000 reads of *swc2*Δ were compared to 1 million reads of WT (**Figure 1D**). SWR-dependent H2A.Z sites were determined by subtracting the normalized distributions of H2A.Z in WT with *swc2*Δ (**Figure 1D**). The normalized H2A.Z read counts at 59,044 annotated nucleosomes (in non-repetitive regions) from two biological replicates were plotted against each other for WT (red) and *swc2Δ* (black) (**Figure 1E**) [39,40]. The data showed strong correlations between replicates (R^2^ = 0.99 for WT; R^2^ = 0.94 for *swc2Δ*) and we identified 10,662 nucleosomal positions with H2A.Z deposited by SWR (*p-values* < 0.05; *null hypothesis:* the H2A.Z signal was noise, which was represented by the *swc2Δ* H2A.Z ChIP-seq data) (**Figure 1E**). Consistent with previous data, SWR deposits H2A.Z predominantly at +1 nucleosomes (3,921 of 4,725 +1 positions) (**Figure 1D**, **1F** and **1G**). However, a substantial amount of H2A.Z was deposited at non-(+1) positions in a SWR-dependent manner (**Figure 1G**). Other NDR-proximal locations, such as +2 and -1 nucleosomes, account for 1,334 and 1,507 H2A.Z-containing nucleosomes, respectively, indicating that there are 3,900 H2A.Z enriched sites that are distal to the NDRs (defined as not +1, -1 or +2) but require SWR for deposition.

In addition, we observed clusters of SWR-dependent H2A.Z sites—termed H2A.Z islands. Using a criterium that defined an island as having ≥ 6 H2A.Z-containing nucleosomes within a 1500-bp window, we identified 264 H2A.Z islands across the yeast genome (**Figure 1D** and **S2A-B**, magenta bars). Many nucleosomes within these H2A.Z islands are not particularly enriched for histone acetylation (**Figure S2C**). In fact, only one third of the NDR-distal H2A.Z sites across the genome are associated with H3 acetylation at lysine 9 (N = 1,510) or H4 acetylation at lysine 12 (N = 1,536) (**Figure 1H**). Therefore, SWR deposits H2A.Z at thousands of distinct H2A.Z sites by an unknown targeting mechanism that is not directly linked to NDRs or histone acetylation.

### SWR preferentially inserts H2A.Z into a subset of nucleosomes normally enriched for H2A.Z in vivo

To identify any unknown determinants for SWR targeting, we developed an unbiased biochemical approach to interrogate the substrate specificity of SWR utilizing a library of nucleosomes prepared from H2A.Z-deficient yeast cells. The latter is important as nucleosomes already inserted with H2A.Z are poor substrates of SWR [26].

To prepare native canonical nucleosomes, we used an *htz1Δswr1Δ* strain that bore an episomal 2xV5-tagged *HHF2* gene as the sole source of H4 histone (**Figure S3**). Logarithmically growing cells were spheroplasted under conditions to minimize proteolytic clipping of histone tails [41]. Nucleosomes and poly-nucleosomes were liberated from the insoluble chromatin pellet by MNase digestion (**Figure 2A-B**). The nucleoproteins were then purified by anti-V5 immunoprecipitation followed by V5 peptide elution. Nucleosomes were separated from poly-nucleosomes by sucrose gradient sedimentation before being dialyzed into a buffer compatible with the remodeling reactions (**Figure 2C** and **S4A**). The native nucleosomal substrate, along with a recombinant nucleosomal control, were incubated with native SWR, ATP and Z-B dimers that were doubly tagged with biotin and FLAG (**Figure 2D, S4B-C**). The native nucleosomes, as well as the recombinant nucleosomes, were active substrates of SWR as indicated by the retardation of the remodeled nucleosomes in native polyacrylamide gel electrophoresis (PAGE) caused by the tags (mainly FLAG) on the Z-B dimer (**Figure 2E**) [28].

**Figure 2.**
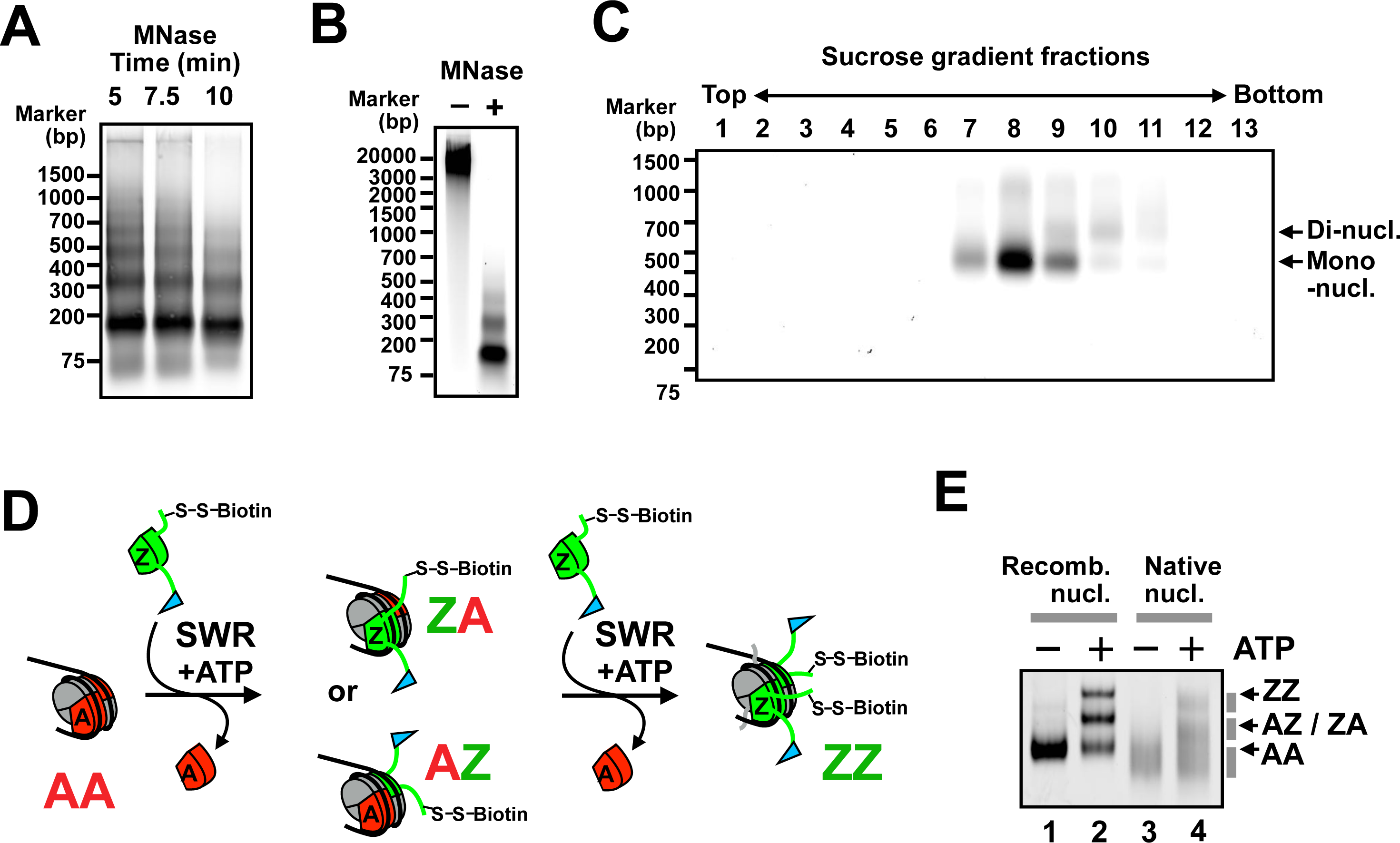
Native nucleosomes isolated from yeast were active substrate of SWR. **(A)** Optimization of MNase digestion. Yeast chromatin was incubated with MNase for the indicated times before extracted for DNA analysis. **(B)** A scale-up MNase digestion before (–) and after (+) MNase treatment. **(C)** Nucleosomes containing V5-tagged H4 were affinity purified by anti-V5 agarose and sedimented through a 15-40% sucrose gradient. The fractions were analyzed by 1.3% agarose / 0.5x TBE electrophoresis and SYBR gold staining. **(D)** The histone exchange assay. Blue flag indicates the 3xFLAG tag on Htb1. The S-S-Biotin label indicates the cleavable biotin tag on Htz1(V126C). Red sectors: A-B dimers. Green sectors: Z-B dimers. Grey sectors: H3-H4 dimers. **(E)** Histone exchange reactions analyzed by 6% polyacrylamide / 0.5x TBE electrophoresis. Recombinant nucleosomes were used in lanes 1 and 2, native nucleosomes in lane 3 and 4.

The nucleosomes remodeled by SWR, i.e., the AZ and ZZ species, were pulled down by streptavidin-coated paramagnetic beads and eluted with dithiothreitol (DTT) as the biotin moiety was linked to H2A.Z (Htz1) via a cleavable disulfide linkage (**Figure 3A**). As a proof of concept, a partially remodeled reaction containing a mixture of recombinant AA, AZ and ZZ nucleosomes (input) were subject to streptavidin pulldown and DTT elution (**Figure 3B** lane 1). The eluate fractions were enriched for the AZ and ZZ species with undetectable AA nucleosomes, indicating that the pulldown was specific (**Figures 3B** lane 3). Similarly native nucleosomes inserted with one or two copies of the dual-tagged Z-B dimers in the SWR-mediated reaction were pulled down by streptavidin beads and liberated into the eluate fraction after DTT treatment (**Figure 3B**, lanes 4-6). To capture the subset of native nucleosomes preferentially remodeled by SWR, we performed reaction time courses. Aliquots of the remodeling reactions, which require Mg^2+^ as a cofactor, were quenched by the addition of ethylenediaminetetraacetic acid (EDTA) at 15, 30, and 45 min (**Figure 3C**). The native nucleosomes were prepared in replicates and reacted with SWR in two independent time courses (**Figure 3C** and **S5A**). The DNA associated with the eluate (EL) and flow-through (FT) fractions of the replicate reactions were purified and analyzed by paired-end sequencing using the Illumina platform (**Figure S5B**).

**Figure 3.**
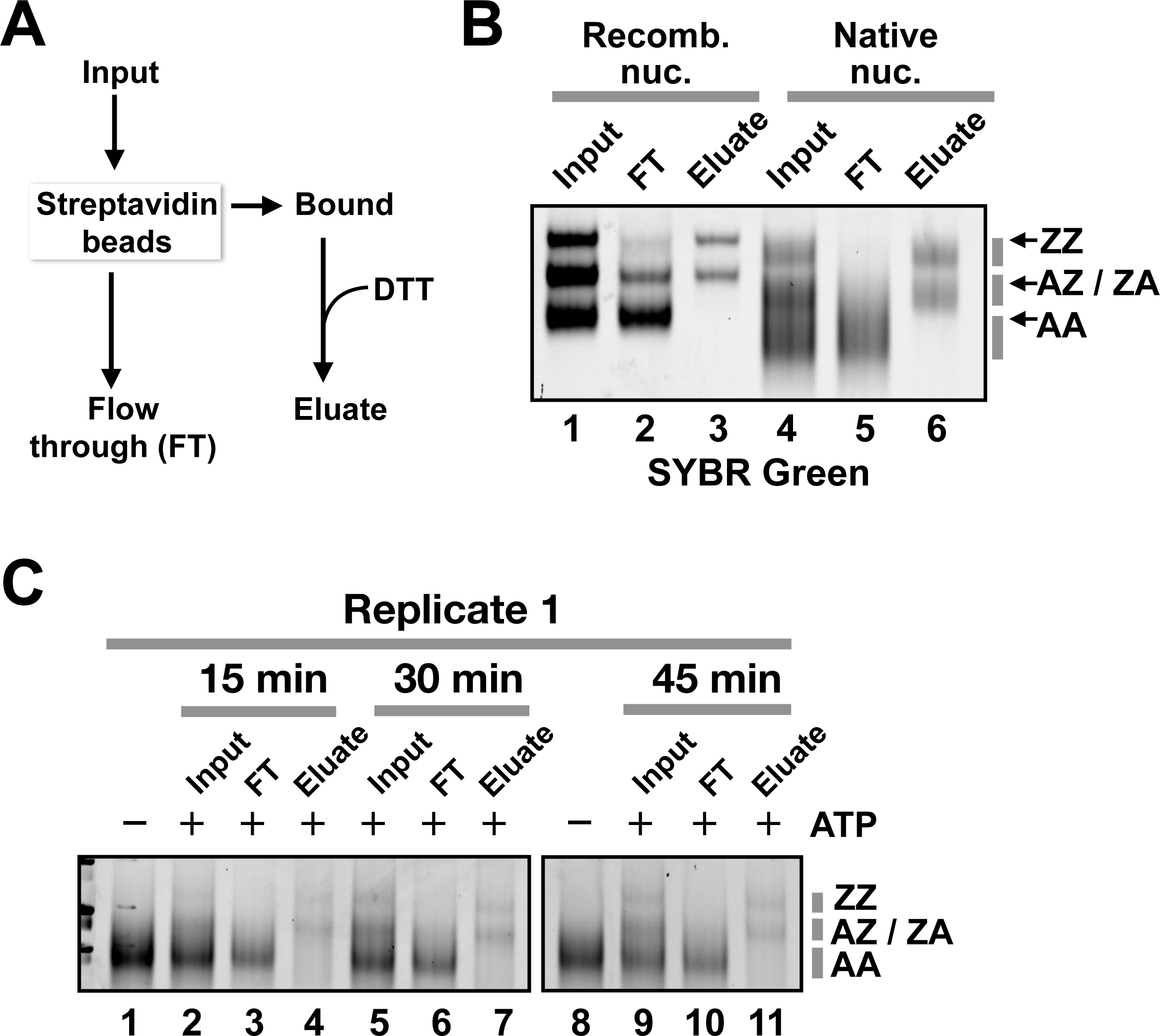
Isolating the biotinylated H2A.Z-containing nucleosomes remodeled by SWR. **(A)** Strategy for isolating SWR-remodeled nucleosomes. Input: nucleosomes partially remodeled by SWR in the presence of Z-B dimers that were biotinylated on Htz1 and FLAG tagged on Htb1. **(B)** Recombinant nucleosomes and native nucleosomes partially remodeled by SWR (input) were pulled down with streptavidin-coated beads and eluted with DTT (eluate). Flow-through fractions (FT) represent the unbound materials of the streptavidin pulldown. Cell equivalent amounts of input, FT, and eluate were analyzed by 6% PAGE and SYBR green staining. **(C)** Same as B except that the remodeling reactions were quenched by EDTA at the indicated times before subjected to streptavidin pulldown.

The sequencing reads of the eluate and FT fractions were mapped to the *S. cerevisiae* genome and the normalized coverages were compared (**Figure 4A**). H2A.Z (Z)-enrichment was defined as the difference of coverage signals in the eluate versus the FT fractions. Although SWR inserted H2A.Z broadly into nucleosomes that cover the bulk of the genome, preferred nucleosomal sites were observed, i.e., when signals of eluate > FT (**Figure 4A**). To evaluate the reproducibility of the site-specific deposition of H2A.Z by SWR, we compared the Z-enrichment values in the two experimental replicates at previously annotated nucleosomal sites (subtracting non-repetitive regions, N = 59,044). The results showed positive correlations of the Z-enrichment values between replicates for all three time points (R^2^: 0.60, 0.52 and 0.51 for 15, 30, and 45 min, respectively) (**Figure 4B**). We defined the top 3% of Z-enriched nucleosomes as the SWR-preferred sites (red) and the bottom 3% as the unpreferred sites (blue) (**Figure 4B**). Importantly, we found the preferred sites were disproportionately associated with nucleosomes that are enriched for H2A.Z in vivo. At the 15-, 30-, and 45-min time points, 32%, 39%, and 44% of preferred nucleosomes were enriched for H2A.Z in vivo, respectively (**Figure 4C**). By contrast, if the sites were picked at random, only 18% of the selected sites were expected to overlap with native H2A.Z nucleosomes (**Figure 1G**).

**Figure 4.**
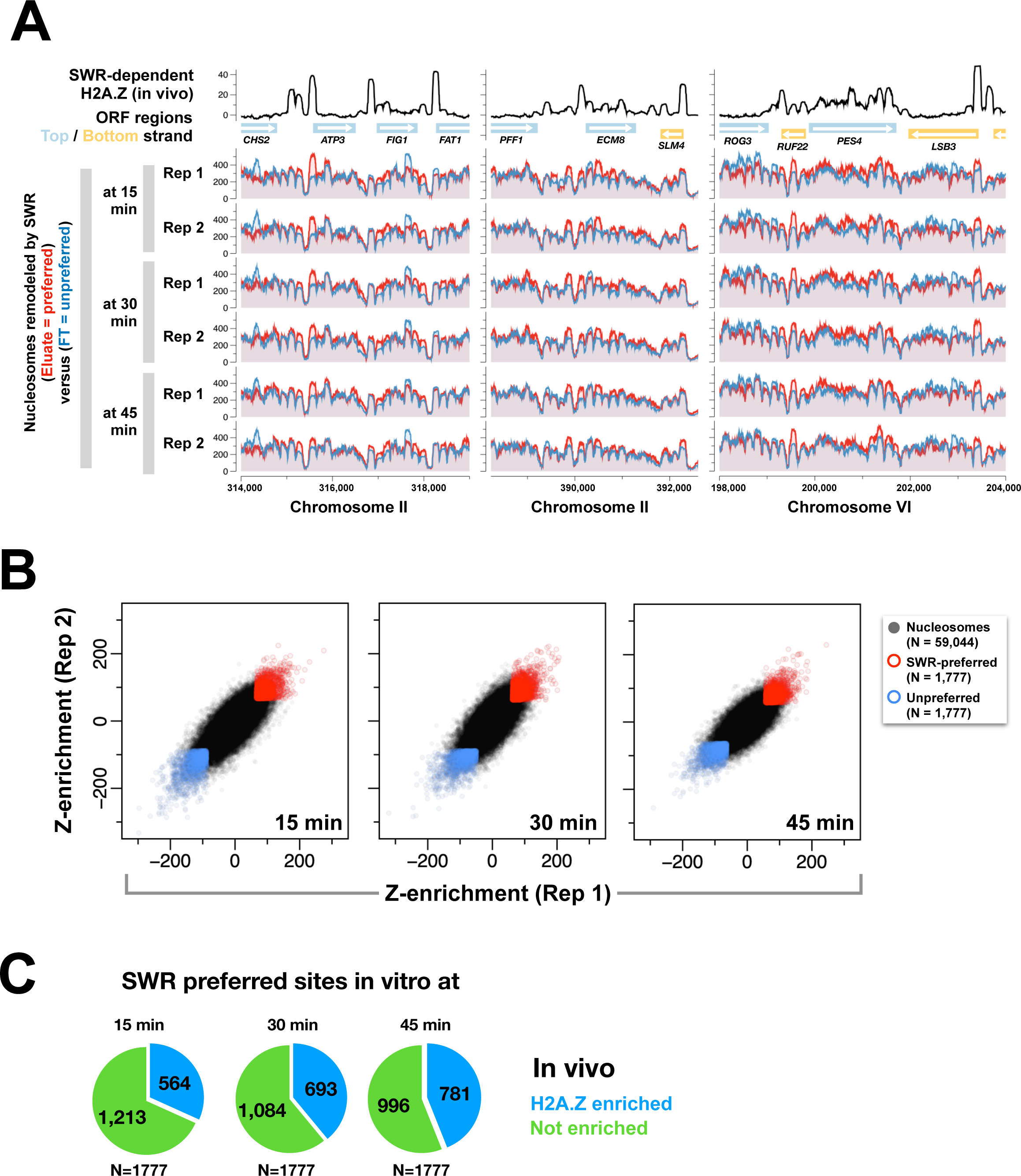
Determination of SWR preferred sites. **(A)** Sequencing analysis of the nucleosomes in the eluate (SWR-preferred) and FT (unpreferred) fractions of the streptavidin pulldown after histone exchange reactions. Red: eluate. Blue: FT. Black traces: endogenous H2A.Z. Three representative regions were shown. **(B)** Z-enrichment values at 59,044 non-repetitive genomic locations for the indicate time points. Red: top 3% of high Z-enrichment values. Blue: bottom 3%. **(C)** Pie charts showing the fraction of SWR-preferred sites (top 3%) that overlap with SWR-dependent H2A.Z sites (blue) versus unenriched sites (green).

### SWR preferentially inserts H2A.Z into nucleosomes associated with intergenic sequences

To further understand how SWR selects its nucleosomal substrates in vitro, we plotted the Z-enrichment values across the yeast genome (**Figure 5A** and **S6**) and aligned the data along the start and end of 4,727 genes (**Figure 5B**). Consistent with the genomic distribution of native H2A.Z, the nucleosomes preferred by SWR in vitro were generally enriched at intergenic regions but depleted across coding regions (compare **Figure 1F** right panel to **5B**). However, SWR exhibited an artifactual preference for the 3’ end of genes not observed in vivo (**Figure 5C**). While SWR-preferred nucleosomes overlap with H2A.Z-enriched +1 nucleosomes, many +1 nucleosomes were missed by SWR in vitro (**Figure S7A**). The bias was particularly prominent at the 15-min time point, while more +1 nucleosomes were inserted with H2A.Z at the 30- and 45-min time points (**Figure S7A**). H2A.Z is enriched at many -1 positions in vivo (**Figure S7A**). Interestingly, SWR has a strong preference for nucleosomes that were slid off-center from the preferred native nucleosomal positions (**Figure S7B**). These off-centered nucleosomes may represent remodeling intermediates during transcription or remodeling events; however, we cannot exclude that they were slid off center artifactually during chromatin extraction.

**Figure 5.**
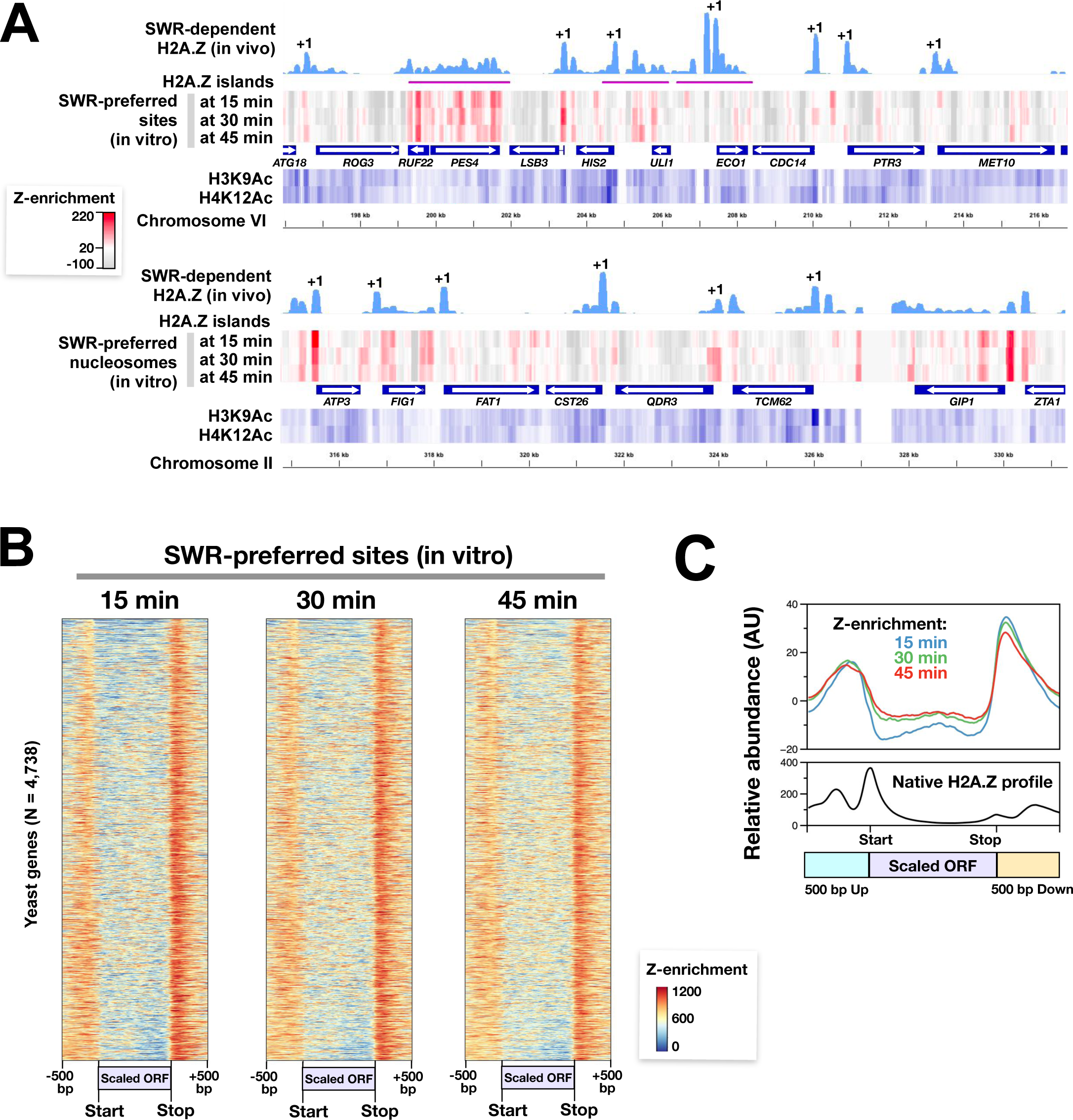
Genomic distribution of SWR-preferred sites. **(A)** Z-enrichment values (red heatmaps) were plotted against the endogenous SWR-dependent H2A.Z (light blue), H3K9Ac and H4K12Ac (blue heatmaps). Two representative regions were shown. Annotated +1 nucleosomes were indicated. **(B)** Heatmaps of Z-enrichment values were sorted using the gene list in Figure 1F right panel. Yeast genes were scaled to equal length and aligned at starts and ends. **(C)** Averaged Z-enrichment plots versus native SWR-dependent H2A.Z (black).

### SWR prefers poly(dA:dT) tracks on the top strand of nucleosomal DNA entry/exit sites

To decipher the sequence composition of the nucleosomal substrates preferred by SWR, we performed dinucleotide motif analysis. Previous studies showed that native nucleosomes in yeast are preferentially positioned on sequences with alternating dinucleotide patterns that appear symmetrical around the nucleosomal dyad and are related to how the two halves of the nucleosomal DNA curved around the histone octamer [43]. We repeated the analysis by plotting the 16 dinucleotide frequencies of 67,534 nucleosome positioning sequences (plus 20 bp linker DNA) (**Figure S8**) [43]. For comparison, the averaged genomic frequencies of the dinucleotides were also shown (**Figure S8**, right panels). In general, dAdA and dTdT dinucleotides are enriched at the minor grooves of the DNA facing the histone octamers, whereas dSdS dinucleotides are out of phase (**Figure S8**). The frequencies of the other dinucleotides (i.e., dAdT, dTdA, dWdS, and dSdW) also alternate in a symmetrical manner around the dyad but their patterns are more nuanced (**Figure S8**).

Having established how dinucleotide frequencies oscillate across positioned nucleosomes, we selected the top 3% of nucleosomes preferred by SWR and asked whether specific dinucleotide motifs are associated with the DNA of the preferred substrate. We found that the preferred nucleosomes were highly enriched for dAdA dinucleotides in the top strand on the left side of the nucleosomal DNA for all three time points, especially in a region between super helical location (SHL) -7.5 and SHL -2.5 and to a lesser extent from SHL -2 to the dyad (**Figure 6A**). The enrichment of dAdA dinucleotides on the right half (top strand) is comparatively less prominent for all three time points (**Figure 6A**). The dAdA dinucleotide pattern was independently observed in the right half of the preferred nucleosomal DNA, which was represented by the enrichment of complementary dTdT dinucleotide pattern in the top strand (**Figure 6B**). Thus, the dAdA and dTdT enrichments observed in the preferred nucleosomes represent a palindromic pattern relative to the nucleosomal dyad. dTdA and dAdT dinucleotides were also more enriched in the nucleosomes preferred by SWR but their enrichments were mainly localized in and around the nucleosomal dyad (between SHL-2.5 and SHL2.5) (**Figure 6C-D**). The increase in dWdW dinucleotide frequencies in SWR’s preferred substrates was accompanied by a decrease in dCdC, dGdG, and dGdC frequencies (**Figure 6E-G**). However, the frequencies of other dinucleotide motifs were largely unchanged in the preferred substrates over the average nucleosome (**Figure 6H** and **S9**). When the dinucleotide frequencies of the unpreferred substrate for SWR (bottom 3%) were compared to those of the averaged nucleosomes, the reverse is generally true, meaning that dWdW dinucleotides were depleted (**Figure S10A-D**), whereas dSdS were enriched in the unpreferred substrates (**Figure S10E-H**).

**Figure 6.**
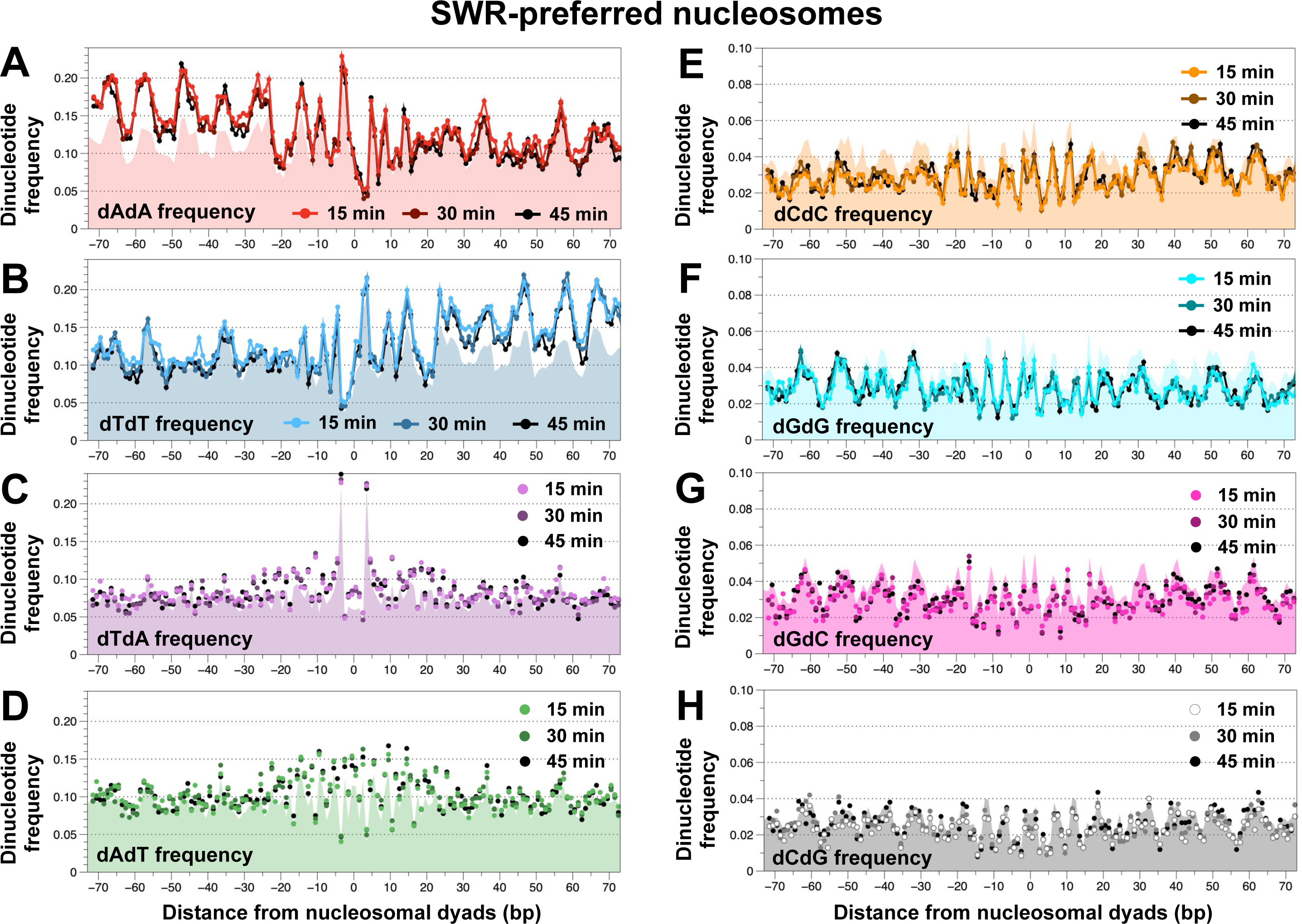
Dinucleotide motif analysis of SWR-preferred nucleosomes. **(A)** The dAdA dinucleotide frequencies of the top 3% SWR-preferred nucleosomes (N = 2,023) for the 15-, 30- and 45-min timepoints were compared to the average (N = 67,534, solid color; replotted using the data from Figure S8). **(B-H)** Same as A, except that the indicated dinucleotide frequencies were shown.

### SWR-mediated H2A.Z insertion activity is stimulated by poly(dA:dT) tracks in nucleosomes

Given that the dAdA and dTdT dinucleotide motifs are enriched in SWR-preferred nucleosomes, we asked if the presence of poly(dA:dT) tracks in the nucleosomal sequence is sufficient to stimulate the H2A.Z insertion activity of SWR in vitro. Using the Widom 601 nucleosome positioning sequence as a platform for mutagenesis, 10 or 13 consecutive dA:dT residues (abbreviated as dA_10_ or dA_13_) were systematically introduced into one half of the nucleosome. The Position (Pos) 1 mutant has a dA_13_ track at the DNA entry site (i.e., between entry site and SHL-6), Pos 2, 3, and 4 have a dA_10_ between SHL-6 and -5, SHL-5 and -4, and SHL-4 and -3, respectively (**Figure 7A**). To ensure that SWR only acts on the half of the nucleosome with the dA tracks, a 2-nt gap was introduced at SWR’s ATPase engagement site on the opposite half of the nucleosome, thereby blocking histone exchange (**Figure 7B**) [28]. To control for the remodeling function of SWR, a reference substrate containing the canonical Widom ‘601’ sequence was simultaneously added to all reactions (**Figure 7C**). Remodeling of both substrates were monitored by fluorescence densitometry on PAGE as the reference and test substrates were end-labeled with Alexa555 and Alexa647, respectively.

**Figure 7.**
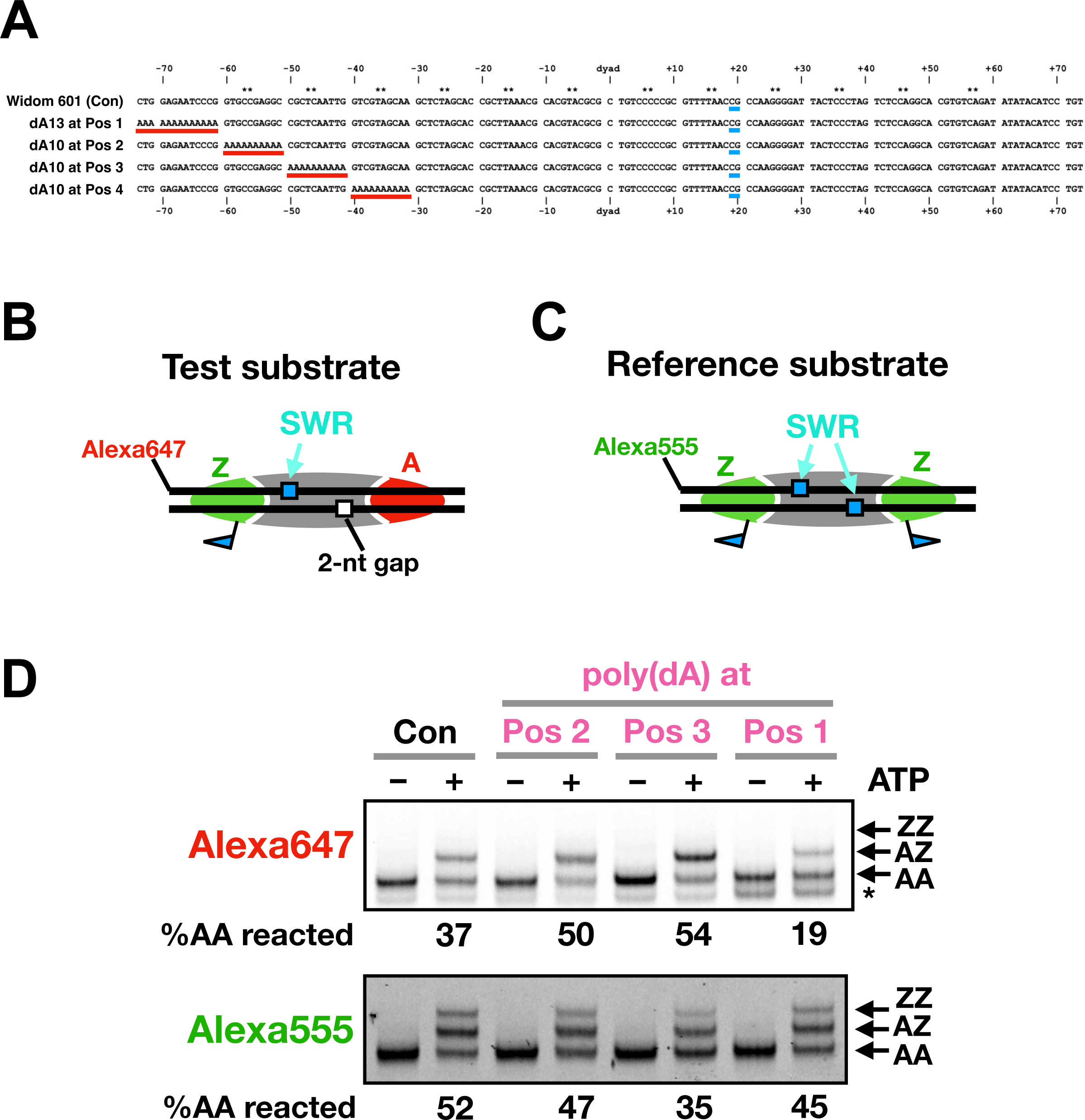
Effects of poly(dA) tracks on H2A.Z deposition. **(A)** Alignment of the Widom 601 control (Con) sequence versus the poly(dA)-containing variants. Red bars highlighted the poly(dA) tracks in positions (Pos) 1-4. Blue bars indicated the position of the 2-nt gap. **(B)** The test substrate has an Alexa647 fluorophore at the 5’ end of the ‘top’ strand and a 2-nt gap at positions +19 to +20 nt (from the dyad) on the ‘bottom’ strand. **(C)** The reference substrate has the canonical Widom 601 sequence and an Alexa555 fluorophore at the 5’ end. **(D)** Fluorescent scans of the nucleosomal DNA. The expected positions for the AA, AZ and ZZ nucleosomes were indicated. Asterisk (*) indicated the expected position for hexasomes. %AA reacted was the difference of AA band intensities in the +ATP lane relative to the no ATP control.

In the control reaction where the test substrate lacks the poly(dA) track, SWR replaced the untagged nucleosomal A-B dimer on the intact side with a FLAG-tagged Z-B dimer, generating AZ nucleosomes which migrated more slowly than AA nucleosomes (**Figure 7D** and **S12A,** top panel). When dA_10_ was inserted at Pos 3, more AZ species were produced relative to the control reaction, indicating that the dA track can indeed stimulate the H2A.Z insertion activity of SWR (**Figure 7D** and **S12A,** top panel). However, the stimulation was position dependent as dA_10_ at Pos 2 stimulated SWR to a lesser extent compared to Pos 3, whereas dA_13_ at Pos 1 and dA_10_ at Pos 4 inhibited SWR’s activity (**Figure 7D** and **S12A-B**). The stimulation observed in the Pos 3 substrate was not due to an artifactual increase of SWR activity in the reaction as the reference substrate in the same reaction was not enhanced but was diminished relative to the control (see bottom panels of **Figure 7D** and **S12**). The reciprocal change suggests that the Pos 3 substrate compete with the reference substrate, further reinforcing the idea that the dA_10_ at Pos 3 makes the nucleosome a better substrate for SWR-mediated H2A.Z deposition.

## DISCUSSION

### A revised model of chromatin remodeling at yeast promoters

The formation of the nucleosome-depleted promoter platform and the closely spaced nucleosome arrays within gene bodies ensure focused assembly of the transcription preinitiation complex and accurate start site selection. The site-specific incorporation of H2A.Z by SWR at the +1 nucleosome of many genes places the variant histone at a juncture whereby RNA Pol II transitions from initiation to elongation, contributing to accurate transcriptional response. In this study, we uncovered an underappreciated role of DNA sequences in H2A.Z deposition, giving new insights into the mechanism on how SWR targets its histone substrate at promoter regions and the sequence of events involved in the biogenesis of the native promoter platform.

Yeast promoters are AT rich with poly(dT) and poly(dA) tracks dominating the sense strand at the upstream and downstream boundaries of the NDR (**Figure 8**, top) [33]. DNA fragments with long duplex poly(dA:dT) are poor substrates for nucleosome assembly as the homopolymeric sequences are rigid and intrinsically curled (i.e., bent and twisted) [44–47]. Thus, yeast promoter sequences are naturally depleted for nucleosomes (**Figure 8**) [48,49]. However, DNA sequence alone is not sufficient to keep ‘nascent nucleosomes’ away from the promoter (deposited presumably after each round of transcription or DNA replication). Chromatin remodeling activities are also required to re-establish and maintain the native NDR observed in vivo [33]. The remodeler RSC, whose activity is stimulated by the poly(dA:dT) tracks, actively clears nucleosomes from the promoter to widen the NDR (**Figure 8** step 2). The remodeler INO80 in turn slides the +1 nucleosome into its native position (**Figure 8** step 3). Where SWR enters this pathway to deposit H2A.Z at the +1 nucleosome is unknown but our data offer some clues.

**Figure 8.**
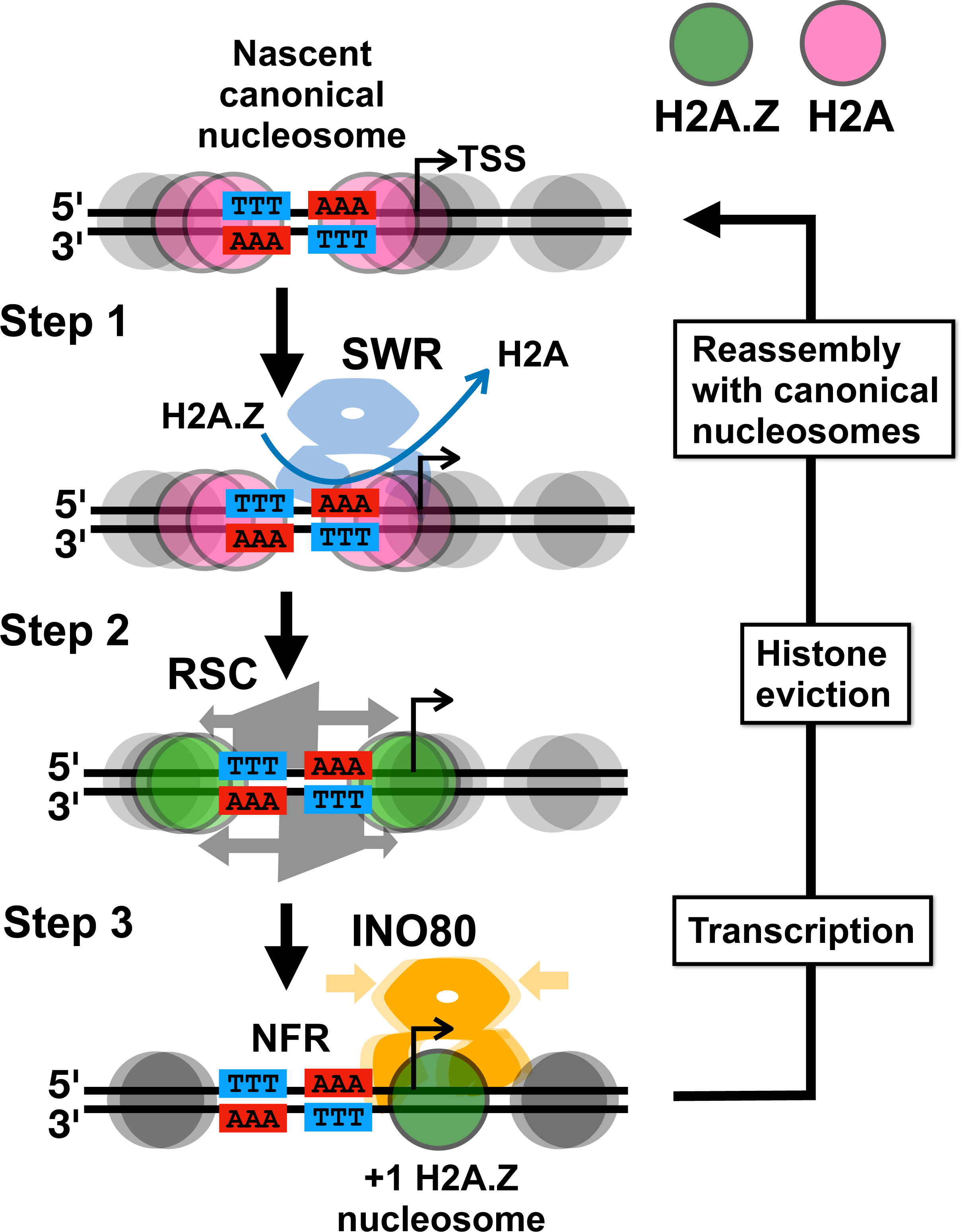
An updated model for the formation of the promoter platform. Red circles: NDR-proximal nucleosomes containing H2A. Green circles: NDR-proximal nucleosomes containing H2A.Z. Grey circles: NDR-distal nucleosomes. AAA and TTT indicates poly(dA) and poly(dT) tracks respectively. Blue crab: SWR complex. Orange crab: INO80 complex. Trapezoid: RSC complex.

We propose that SWR acts upstream of RSC and INO80 to replace A-B dimers in nascent, promoter-proximal nucleosomes with Z-B dimers. The rationale is three-fold. First, before a nascent nucleosome is sled into its native +1 position by RSC and INO80, its upstream edge overlaps with the poly(dA:dT) sequence, hence placing the poly(dA) track on the top strand of the nucleosomal DNA between SHL-4 and -6. This will make the nascent nucleosome a preferred substrate for SWR-mediated histone exchange. Second, promoter-proximal deposition of H2A.Z is observed in yeast conditionally depleted for RSC (Sth1) [34]. This suggests that positioning of the nucleosome at the native +1 position is not a prerequisite for SWR-mediated histone exchange. Third, INO80 preferentially slides H2A.Z-over H2A-containing nucleosomes [50]. Therefore, it is energetically more efficient to have SWR deposited H2A.Z before INO80 slides the +1 nucleosome into its native position (**Figure 8**).

### How poly(dA:dT) sequences facilitate H2A.Z insertion

When SWR was presented with native nucleosomes isolated from H2A.Z-depleted cells, Z-B dimers were preferentially deposited into the subpopulation of nucleosomes associated with intergenic regions over genic regions. While extra precaution was taken to enrich mono-nucleosomes by sucrose gradient sedimentation, we cannot rule out that the preferred nucleosomes may be bound by non-histone factors that indirectly contribute to the recruitment of SWR. Notwithstanding this potential caveat, the overrepresentation of poly(dA:dT) sequences in the entry sites and the A-B dimer contact sites of the preferred nucleosomes is clear. At the mechanistic level, the DNA sequences may introduce structural stress such that the entry DNA can ‘spring open’ more easily as the Swr1 ATPase at SHL-2 draws in DNA to dissociate the histone-DNA contacts of the outgoing A-B dimer, thereby facilitating the coordinated Z-B dimer insertion. Our in vitro experiment is consistent with this model as the poly(dA) tracks only stimulated H2A.Z insertion when they are present at SHL -4 to -6, which overlaps with the contact site of the outgoing A-B dimer. Why SWR prefers poly(dA:dT) tracks but not poly(dT:dA) tracks is intriguing. This suggests that poly(dA:dT) may curl the entry DNA in such a way that is more compatible with the transition state of SWR-mediated histone exchange, whereas the curl caused by poly(dT:dA) is reversed and is not particularly favorable.

### Intrinsic bias of SWR against genic nucleosomes

SWR preferentially deposits H2A.Z into nucleosomes associated with intergenic over genic sequences. One explanation is that the information-rich genic sequences are depleted for the monopolymeric poly(dA:dT) sequences preferred by SWR. In fact, when genic regions were subject to dinucleotide motif analysis, 3-bp periodicities were observed for all 16 dinucleotide motifs, revealing the biased codon usage and codon-pair preference of yeast genes (**Figure S13**). Therefore, poly(dA:dT) sequences are selected against in the coding region, hence unpreferred by SWR. An exception is with the genes that encode lysine-rich and phenylalanine-rich domains, i.e., with consecutive AAA and TTT codons (**Figure S14**). This provides an explanation for why some genes in the H2A.Z islands, such as *PES4* (**Figure 5A**), exhibit non-canonical H2A.Z deposition in genic regions.

Finally, SWR exhibits a strong preference for nucleosomes at the 3’ end of genes, consistent with the presence of poly(dA) tracks (on the sense strand) associated with the transcriptional termination signal in yeast. However, unlike the preferred H2A.Z insertion at the promoter regions, the preferred insertion at 3’ end is non-physiological, as native H2A.Z are generally enriched at the 5’ end but not the 3’ end of genes. How poly(dA:dT) sequences at terminator sequences exclude SWR-mediated H2A.Z insertion in cells is unknown and may involve masking by termination factors in the regions [34].

In conclusion, this study provides direct evidence showing that the context of nucleosomal DNA sequences can influence the H2A.Z insertion activity of the SWR complex. Thus, SWR is now added to a growing list of chromatin remodelers, including RSC and Chd1, whereby their remodeling activities are tuned by DNA sequences [37,42]. One emerging idea is that the genome may be encoded with information that can direct chromatin remodeling activities and/or that remodelers are evolved to recognize sequence signatures associated with specific genomic regions, such as promoters [35]. This study also underscores the power of native yeast nucleosomes as substrates to interrogate chromatin remodeling events. This biochemical approach when combined with high-throughput sequencing analysis can reveal how substrate diversity can affect chromatin remodeling events. Although the preferred nucleosomes of SWR under our reaction conditions do not appear to be linked to known post-translational modifications, our approach can in theory be used in conjunction with mass spectrometry to identify other histone marks preferred by remodelers in an unbiased manner, paving ways for future studies.

## Materials and Methods

### Yeast strains

The WT strain (yEL379) in the VivosX analysis of Figure 1B-C was previously described in [38]. The *swc2Δ* strain (yEL575) was constructed by replacing the *SWC2* ORF in yEL379 with the *kanMX6* cassette using the homologous gene replacement method [43,44]. The WT strain (yEL378) in the ChIP-seq analysis was previously reported [38]. The corresponding *swc2*Δ variant (yEL786) was similar constructed by homologous gene replacement.

The yeast strain (yEL905), which was used to prepare native nucleosome libraries, carried an episomal *HHF2* gene with a 2xV5 tag at the N-terminus to facilitate nucleosome isolation. In addition, the strain lacked *HTZ1* and *SWR1* to remove endogenous H2A.Z. To construct yEL905, the precursor strain YYY67 was used [45]. YYY67 lacked the endogenous H3 and H4 genes, i.e., (*hht1-hhf1*)Δ::*LEU2* (*hht2-hhf2*)*Δ*::*HIS3* and was kept alive by the an *HHT1-HHF1 URA3 CEN ARS* (pMS329) plasmid. A *TRP1 CEN ARS* plasmid (pEL628) containing the divergently oriented *HHT1* and 2xV5-*HHF2* genes was transformed into YYY67 and was used to replace pMS329 based on the plasmid shuffle technique [46]. The resulting strain (yEL704), which expressed the 2xV5-*HHF2* gene as the sole source of H4, was subjected to a second round of transformation to replace *HTZ1* with the *hphMX*6 cassette [43]. Finally, the strain was subjected to a third round of transformation to replace *SWR1* with *kanMX6*. Successful removal of the *HTZ1* and *SWR1* genes was verified by colony PCR.

### Plasmids

The *2xV5-HHF2* gene for native nucleosome isolation was subcloned into a *HHT2-HHF2 TRP1 CEN ARS* plasmid (gift from Rolf Sternglanz [45]) at the *HHF2* location by digestion with BamHI and NcoI followed by recombination with a synthetic *2xV5-HHF2* fragment using the Gibson Assembly protocol (New England Biolabs).

The plasmid for glucanase production (pT22-6HisLyticase) was constructed as follows. The sequence encoding amino acid residues 37-548 of glucanase (glucan endo-1,3-beta-glucosidase from *Cellulosimicrobium cellulans*, Uniprot: P22222) was synthesized by IDT and amplified by PCR using primer pT22(NdeI)Gluc-Gibs-F and pT22(XhoI)Gluc-Gibs-R (**Table S1** and **S2**). The PCR product was cloned into the pET22b vector (Novagen) pre-digested with the NdeI and XhoI restriction enzymes (Thermo) using the HiFi Gibson Assembly mix (NEB). The sequence integrity of all plasmids was confirmed by Sanger Sequencing (Azenta).

### VivosX

In vivo crosslinking analysis was performed according to a previous study [38]. Briefly, cells were grown in the complete synthetic media (CSM) to an optical density at 600 nm (OD_600_) of 0.5 before 5-mL aliquots were treated with 180 uM 4-DPS (Sigma; Cat# 143057-1G) for 20 min at 30°C. The cells were fixed on ice by adding 20% trichloroacetic acid (TCA) for > 5 min. Fixed cells were collected by centrifugation at 2,851 xg for 5 min and washed once with 20% TCA. The cells were lysed in 400 µL 20% TCA with ∼400 µL 0.7-mm zirconia beads using a FastPrep-24 homogenizer (2x 30 sec at 6.0 M/s speed). The insoluble material contained the crosslinked histones and were collected by centrifugation at 20,400xg for 15 min at 4°C. The pellet was washed with 1 mL acetone. To extract proteins from the pellet, 200 uL of the TUNES-G Buffer [100 mM Tris pH 7.2, 6 M urea, 10 mM EDTA, 1% SDS, 0.4 M NaCl, 10% glycerol] + 50 mM N-ethylmalemide (Sigma; Cat# E1271-5G) was added and the dispersed precipitates were vortexed for 1 hr at 30°C. The solubilized proteins were collected by centrifugation at 20,400xg for 10 min at 4°C and analyzed on a non-reducing 8-16% polyacrylamide gel. Immunoblotting analysis was performed by transferring the proteins to a PVDF membrane and probing with the anti-FLAG or anti-H4 primary antibodies (at 1:1000) and anti-mouse HRP (at 1:2000). Immunoblots were developed using the Amersham ECL Prime detection reagent (Cytiva; Cat# 45001216) and imaged on a LAS4010 CCD camera (GE Healthcare).

### ChIP-seq

Chromatin was extracted from the yeast strains yEL378 (*SWC2 HTZ1-2FLAG*) and yEL786 (*swc2Δ HTZ1-2FLAG*) as described with minor modifications [16]. Briefly, cells (uncrosslinked) equivalent to a 400-mL culture at 0.5 OD_600_ was spheroplasted and lysed in 1x pellet volume of Buffer A [50 mM HEPES (pH 7.6), 80 mM NaCl, 0.25% Triton X-100, with 2x Complete Protease inhibitors (EDTA-free, Roche; Cat# 5056489001)] using a Dounce homogenizer. After centrifugation, the chromatin-enriched pellet was washed three times with Buffer A before it was resuspended in Buffer A supplemented with 1 mM CaCl_2_. The crude chromatin suspension was digested with 1 U/µL MNase (Worthington; Cat# NC9391488) for 20 min. The solubilized nucleoproteins were cleared by centrifugation at 20,400xg for 5 min at 4°C followed by filtration using an Ultrafree-MC spin column (Millipore, UFC30GV0S).

ChIP-seq was performed as previous described but with some modifications [16]. The solubilized nucleoproteins were diluted 10 folds in Buffer B [25 mM HEPES (pH 7.6), 1 mM EDTA, 80 mM KCl, 1x protease inhibitor cocktail (PI)] before incubated with 200 µL (slurry volume) of anti-FLAG M2 affinity gel (Sigma; Cat# A2220) per 5-mL diluted nucleoproteins. Binding was performed at 4°C for 4 hr on a rotator. The FLAG beads were washed with Buffer C [25 mM HEPES (pH 7.6), 1 mM EDTA, 0.01% NP-40, 0.3 M KCl, 1x PI] and eluted with 0.5 µg/µL 3xFLAG pepetides (Biopeptide) in the Buffer D [HEN_0.1%_-0.3_K_-Doc_0.1%_-P [25mM HEPES (pH 7.6), 1mM EDTA, 0.1% NP-40, 0.3M KCl, 0.1% DOC, 1x PI] overnight at 4°C. Nucleosomal DNA was extracted by incubating with 0.34 µg/µL Proteinase K (Invitrogen; Cat# 25530049) in the Buffer E [340 mM NaCl, 8.5 mM EDTA, 0.43% SDS, 0.1 ug/uL Glycogen] at 55°C for 1 hr followed by extraction with phenol:chloroform:IAA (25:24:1) and ethanol/sodium acetate precipitation. The nucleic acid pellet was resuspended TE buffer and treated with 0.05 µg/µL RNase (Roche Cat# 11119915001) for >15 hr at 37°C. The DNA was purified using the QIAquick PCR purification kit and it was prepared for Illumina sequencing using the NEBNext Ultra II A Library Prep Kit (New England Biolabs; Cat# E7645L).

### Statistics

To identify SWR-dependent H2A.Z nucleosomes, normalized H2A.Z read counts were computed at annotated nucleosomes within non-repetitive regions (N = 59,044) for both WT and swc2 samples. The difference in H2A.Z signals between WT and swc2 represents SWR-dependent H2A.Z. Using a predefined threshold of 10 (i.e., WT-swc2 >= 10 for both replicates), we identified a total of 10,662 SWR-dependent nucleosomal H2A.Z sites. To assess the significance of H2A.Z signals at these sites, a statistical test was performed, positing the null hypothesis that these signals represent background noise. Utilizing the normally distributed H2A.Z ChIP-seq data from the *swc2Δ* strain as a noise model, Z-scores and the associated p-values were computed under the assumption of the null hypothesis. The maximum p-value observed among the 10,662 H2A.Z sites in WT replicate 1 was 0.015, and in WT replicate 2, it was 0.028. These outcomes led to the rejection of the null hypothesis, indicating that the 10,662 SWR-dependent H2A.Z sites exhibit a significant enrichment of H2A.Z.

### Proteins

#### Yeast SWR complex

Native SWR was purified from total extracts from a yeast strain (yEL427) bearing the *SWR1-3xFLAG* and *RVB1-MBP* alleles using the Another Sequential Affinity Purification protocol previously described [28]. The concentration of SWR was quantified by in-gel staining using the SYPRO Orange dye against a known concentration gradient of bovine serum albumin protein (Roche).

### Recombinant histone substrates

The dual tagged Z-B dimer has a cleavable biotin tag conjugated to a cysteine residue substituted at valine position 126 (V126C) of the yeast Htz1 and a triple FLAG tag at the C-terminus of yeast Htb1 [26,28]. Both proteins were produced recombinantly using the *E. coli* strain BL21 Codon Plus (DE3)-RIL and purified according to a previous protocol [47]. Biotinylation of Htz1(V126C) was carried out using N-[6-(Biotinamido)hexyl]-3’-(2’-pyridyldithio)propionamide (HPDP-Biotin) (Thermo Fisher cat. # 21341), which has a pyridyl disulfide moiety. To ensure optimal biotinylation, 48 µM of Htz1(V126C) protein was first dissolved in Buffer F [25 mM HEPES (pH 7.6), 10 mM EDTA, and 0.15 M NaCl] and pre-treated with 5 mM DTT at 37°C for 1 hr. Excess DTT was removed by gel filtration using a PD-10 column equilibrated with Buffer F. The reduced Htz1(V126C) (in ∼3.5 mL) was concentrated to 1 mL on an Amicon Ultra-4 (10kD MWCO) column (Millipore) before 100 µL of 4 mM HPDP-Biotin [in dimethylformamide (DMF)] was added to allow disulfide exchange at 20.8°C for 1.5 hr. To remove any unreacted HPDP-Biotin, the reaction was applied to a new PD-10 column and eluted with 3.5 mL Buffer F. The eluted protein was then unfolded by the addition of 5.02 g of guanidine-HCl and 50 µL 1M Tris-HCl (pH 7.5). Molecular grade water was added to a final volume of 7.5 mL before 2 mg of Htb1-3xFLAG protein was mixed with the biotinylated Htz1 to unfold at 20.8°C for 2 hr. Protein refolding was performed by dialysis against four changes of 2 L of Buffer G [10 mM Tris-HCl (pH 7.5), 2 M NaCl, 1 mM EDTA, 0.2 mM PMSF]. The refolded dimers were concentrated to ∼500 µL (using Amicon Ultra-4) and purified on a Superdex 200 Increase 10/300 GL column (Cytiva) equilibrated with Buffer G. The peak fraction was dialyzed into Buffer H [10 mM Tris-HCl (pH 7.5), 50 mM NaCl, 1 mM EDTA, 0.01% NP-40] and stored in aliquots at -80°C before ready to use in nucleosome remodeling reactions.

Recombinant nucleosomes were made by combining canonical yeast histone octamers with PCR synthesized DNA followed by refolding using the salt gradient dialysis approach previously described [28]. Recombinant yeast H2A and H2B histones were purchased from the Histone Source (CSU Fort Collins, CO) and yeast H3 and H4 were produced as described [28]. The primers and templates used to generate the Widom 601 containing fragments and the poly(dA)-containing variants are listed in **Tables S1-S2**. The amino-modified forward primers (synthesized by IDT) were chloroform extracted twice before labeling with Alexa647- or Alexa555-conjugated ester according to the manufacturer protocols (Thermo Fisher, A20106 and A20109). The labeled primers were purified by PAGE and were used in combination with reverse primer EL338 or EL873 to amplify eBlock gene fragments (IDT), gEL172-gEL176, with or without the poly(dA) sequences. PCRs (6.076 mL each) were performed in 96-well plates using *taq* polymerase. PCR products were purified by phenol extraction and concentrated by ethanol precipitation as described above before injected into a Superose 6 Increase 30/100 GL column equilibrated with Buffer I [10 mM Tris-HCl (pH 7.5), 1 mM EDTA, 300 mM NaCl]. Fractions containing the labeled PCR products were concentrated to ∼400 µL using Amicon Ultra 4 spin columns. To introduce the 2-nt gap in the dUdU modified DNA, 100 µg of DNA was incubated with 0.1 U/µL of USER enzyme (NEB) in 500 µL at 37°C overnight. The DNA products were purified by phenol extraction and concentrated by EtOH precipitation.

### Glucanase

Production of recombinant glucanase was performed using the *E. coli* strain BL21 DE3 RIPL transformed with the pT22-6HisLyticase plasmid. The cells were grown in 6x 500 mL of TB supplemented with 100 µg/mL of ampicillin to log phase at OD_600_ 0.5. Glucanase production was induced with 0.5 mM IPTG and was allowed to proceed at 25°C overnight. The cells were harvested by centrifugation, resuspended in Buffer J [50 mM Tris-HCL (pH7.6), 10 mM Imidazole, 500 mM NaCl, 3 mM beta-mecaptoethanol, and 0.2 mM PMSF] and sonicated using a microtip on a Q Sonica sonicator at 50% power for 3 min (15 sec ON, 30 sec OFF). The supernatant was clarified by centrifugation and applied to a 5-mL HisTrap HP column and eluted with a 10-400 mM linear imidazole gradient on an AKTA Pure FPLC system (Cytiva). Peak fractions were dialyzed into Buffer K (2 mM MES pH 6, 10% sucrose, 10% glycerol, 5 mM DTT) and loaded onto a 5-mL SP HP column. The protein was eluted with a linear gradient of 0-750 mM NaCl in Buffer K and analyzed by SDS-PAGE, concentrated 8-fold to 500 µL, flash frozen, and stored at -80°C.

### Native nucleosomes

Native canonical nucleosomes were prepared from the yeast strain yEL905 that lacked the endogenous *HTZ1*, *SWR1*, *HHT1-HHF1,* and *HHT2-HHF2* genes and expressed an N-terminal 2xV5-tagged H4 from an *HHT2-(2xV5-HHF2) TRP1 CEN ARS* plasmid. Logarithmically growing cells (at 0.5 OD_600_) cultured in 2 L yeast extract peptone dextrose (YPD) media were harvested by centrifugation, washed, aliquoted, and flash frozen. Each cell pellet equivalent to a 400-mL of culture was thawed and washed with 6 mL of Buffer L [100 mM Tris (pH 9.4), 10 mM DTT] and incubated for 5 min at 30 min in 4 mL of Buffer M [50 mM KPO_4_ (pH 7.5), 0.6 M Sorbitol, 10 mM DTT] plus Inhibitor Cocktail (InhC) [1 mM sodium fluoride, 10 mM beta-glycerophosphate, 0.5 µM trichostatin A, and 10 mM sodium butyrate and 2x Complete EDTA-free protease inhibitor mix (Roche)]. The cells were spheroplasted to 80% completion by adding 40 µL glucanase and incubated at 30°C for ∼15 min. Spheroplasts were washed three times with 8 mL Buffer N [50 mM HEPES (pH 7.5), 80 mM KCl, 2.5 mM MgCl_2_, 0.4 M Sorbitol, and InhC] resuspended in 500 µL Buffer O (same as Buffer A except that the protease inhibitor mix was replaced with InhC). The spheroplasts were disrupted in a pre-chilled 7-mL Dounce homogenizer with a tight piston at 4°C. The crude chromatin was pelleted by centrifugation at 13,000 x g for 10 min, washed three times with 500 µL Buffer O and resuspended in 500 µL Buffer O. Chromatin fragmentation was initiated by adding 0.1 mM CaCl_2_ and 0.3 U/µL MNase. The digestion was allowed to proceed at 37°C for 5 min followed by quenching with 10 mM EDTA. The supernatant (chromatin solution), which contained the liberated nucleoprotein fragments, was clarified by centrifugation at 20,400xg for 5 min and filtration using an Ultrafree-MC spin column (Millipore, UFC30GV0S).

To pulldown nucleoproteins containing the 2xV5-H4 histone, the chromatin solution from the two 400-mL equivalent of cultures were pooled (800 µL total) and diluted to 8 mL using Buffer P (same as Buffer B except that the protease inhibitor mix was replaced by InhC) and incubated with 200 µL (slurry volume) of anti-V5 beads (Sigma) at 4°C for 4 hr. Elution was performed using Buffer Q [25mM HEPES (pH 7.6), 0.3M KCl, 1mM EDTA, 0.1% NP-40, 0.1% DOC, and 0.5 µg/µL V5 peptides] at 4°C overnight. The eluate was concentrated on a Amicon Ultra-4 centrifugal filter unit and sedimented through a 4.7-mL15-40% sucrose gradient in 25 mM HEPES (pH7.6), 0.5 mM EDTA, 0.01% NP-40 at 45,000 RPM (366,613 xg) for 20 hr at 4°C using a Beckman SW55Ti rotor. The gradient was fractioned by pipetting and analyzed by electrophoresis in 1.3% agarose/0.5x TBE and SYBR Green I staining. Fraction 8 (top fraction is 1) represent the peak fraction of mononucleosomes and was collected and dialyzed against two changes of 1 L Buffer H.

### In vitro assays

Histone exchange reactions were performed by mixing the following components. For a 25-µL reaction, 11.4 µL of Buffer R [25 mM HEPES-KOH (pH 7.6), 0.5 mM EGTA, 0.1 mM EDTA, 5 mM MgCl_2_, 0.17 μg/μL BSA, 50 mM NaCl, 10% glycerol and 0.02% NP-40] was mixed with the biotinylated, FLAG-tagged Z-B dimers (to 25 nM in 25 µL) in a 1.5-mL LoBind tube (Eppendorf). Then canonical nucleosomes (from native or recombinant source) were added (to 29 nM in 25 µL) followed by the addition of SWR (to 1.5 nM in 25 µL). Buffer H was added to adjust the reaction volume to 20 µL. The reaction was initiated by adding 5 µL of 1 mM ATP in Buffer S [25 mM HEPES-KOH (pH 7.6), 0.5 mM EGTA, 0.1 mM EDTA, 0.02% NP-40] or 5 µL of Buffer S (no ATP control). The reactions were allowed to proceed at room temperature for the indicated times and were quenched by adding 5 µL of 10 mM EDTA. For native PAGE analysis, 5 µL of the reaction was mixed with 1 µL of Buffer T [70.5% w/v sucrose in 10 mM Tris-HCl (pH 7.8), 1 mM EDTA] plus 1 µL 20 µg/µL lambda phage DNA (NEB, cat # N3011S) and was separated on a 6% polyacrylamide / 0.5x TBE / 1.5 mm gel for 2 hr at 110 V in a Novex electrophoresis system. The gel was stained with SYBR Green I (Thermo) and imaged using the Typhoon FLA9500 scanner (Cytiva).

To isolate the biotinylated product of SWR, histone exchange reactions were scaled up to 100 µL for each pull down reaction. Streptavidin-coated M-280 Dynabeads (Thermo Fisher; cat. #112.50) (20 µL slurry volume) prewashed with Buffer H in a 1.5-mL LoBind tube (Eppendorf) were incubated with 80 µL of the reaction. The remaining 20 µL of the reaction was set aside as the ‘input’. The mixture was incubated at room temperature for 30 min with gentle ‘flicking’ of the tube every 5 min. The streptavidin-coated beads were pulled down with a magnet. The supernatant was set aside as the FT fraction. The beads were washed once with 500 µL of Buffer H and eluted with 80 µL of Buffer H supplemented with 1 mM DTT at room temperature for 30 min with gentle flicking every 5 min. Five microliters of the input, FT, and eluate fractions were mixed with 1 µL of Buffer T and analyzed by PAGE and SYBR Green staining as described above. The remaining sample of the flow-through and Eluate fractions were analyzed by next-generation sequencing using the Illumina platform.

To purify the nucleosomal DNA for sequencing, the FT and Eluate fractions (90 µL each) were mixed with 10 µL 3 M sodium acetate and applied to a QIAquick spin column (Qiagen). Purified DNA was visualized a 1.3% agarose / 0.5x TBE gel and SYBR Green staining. DNA concentrations were determined using the Qubit High Sensitivity (HS) double-stranded DNA assay (Invitrogen) on a Biotek Synergy 2 plate reader.

### Sequencing and bioinformatics

Sequencing libraries were prepared using the NEBNext Ultra II DNA library prep kit according to the manufacturer protocol (NEB). Sequencing libraries were pooled and analyzed on a NextSeq 550 platform using paired-end mode with 32 cycles on both ends. Sequencing data (FASTQ files) were mapped to the *S. cerevisiae* S288C genome version R64-1-1 using Bowtie2 with options *--local --very-sensitive-local --no-unal --no-mixed --no-discordant --phred33 -I 120 - X 170*. Mapped reads were filtered using Samtools-view with options *-q 30 -F 0×4*. Read coverage were calculated using bamCoverage in deepTools with options *-binSize 1 -- extendReads --effectiveGenomeSize 11756462 --scaleFactor X* (Supplemental file 1) -- *blackListFileName* Mask_v2.bed (which indicated repetitive sequences in the BED format) (Supplemental file 2).

## Acknowledgements

This work was supported by the NIH grants R01GM104111 and R01GM147795 to E.L. and R35 GM149279 to J.M.D. and the NIH Shared Instrumentation Grant (SIG) S10OD024986 to the Genomics Core Facility of Stony Brook University.

## Supplemental Figure Legends

**Figure S1.**
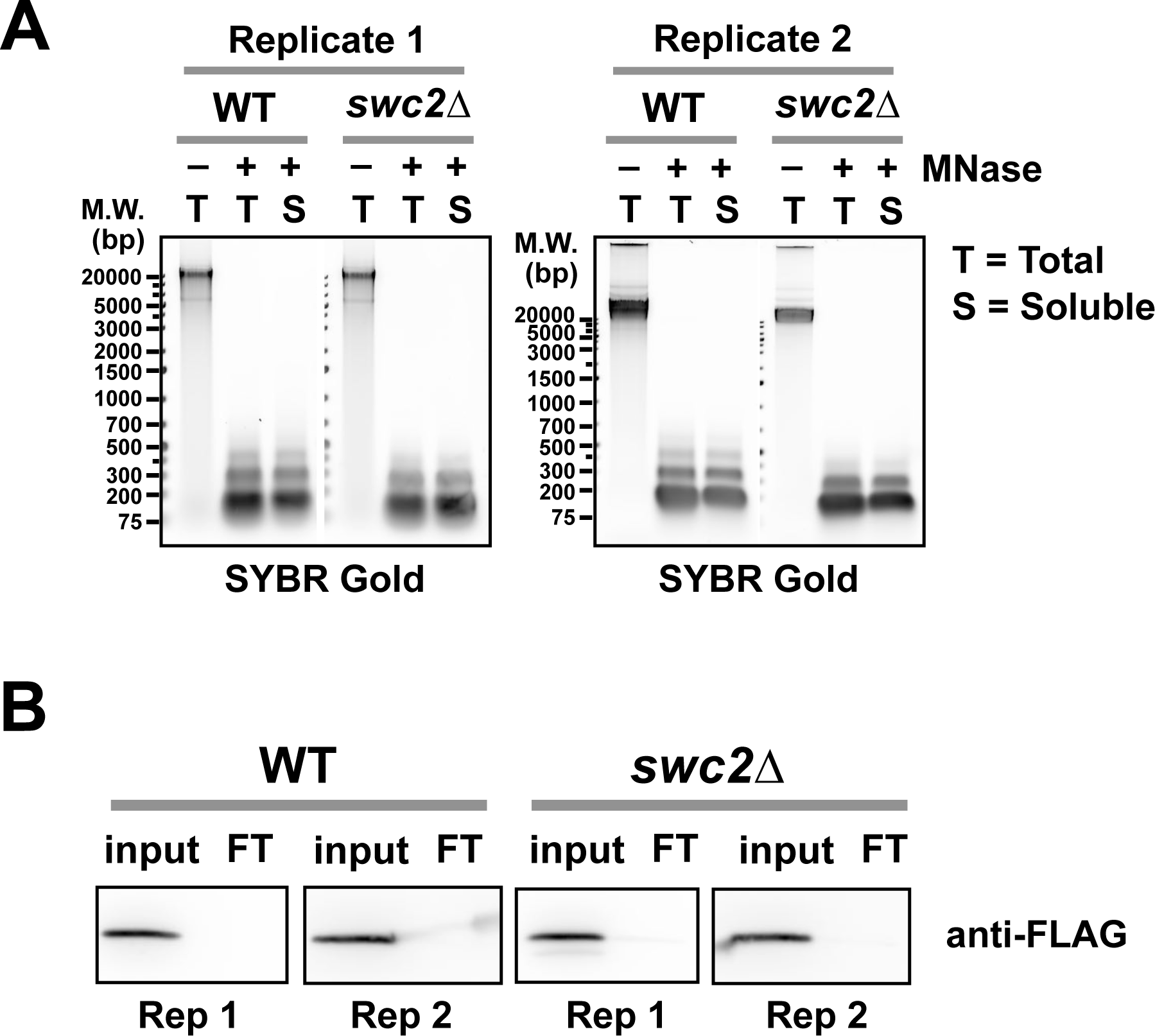
Quality control of the samples used for the H2A.Z ChIP-seq analysis. **(A)** Control for MNase digestion. Chromatin before (-) and after (+) MNase treatment from WT and *swc2Δ* cells were analyzed by agarose gel electrophoresis and SYBR Gold staining. T: DNA extracted from total extracts. S: DNA extracted from soluble fractions, which were used in the anti-FLAG pulldown reactions against Htz1-2xFLAG. **(B)** Control for IP efficiency. Equivalent amounts of soluble chromatin before (input) and after immunoprecipitation (FT or flow through) were analyzed by SDS-PAGE and anti-FLAG immunoblotting. Rep: replicate.

**Figure S2.**
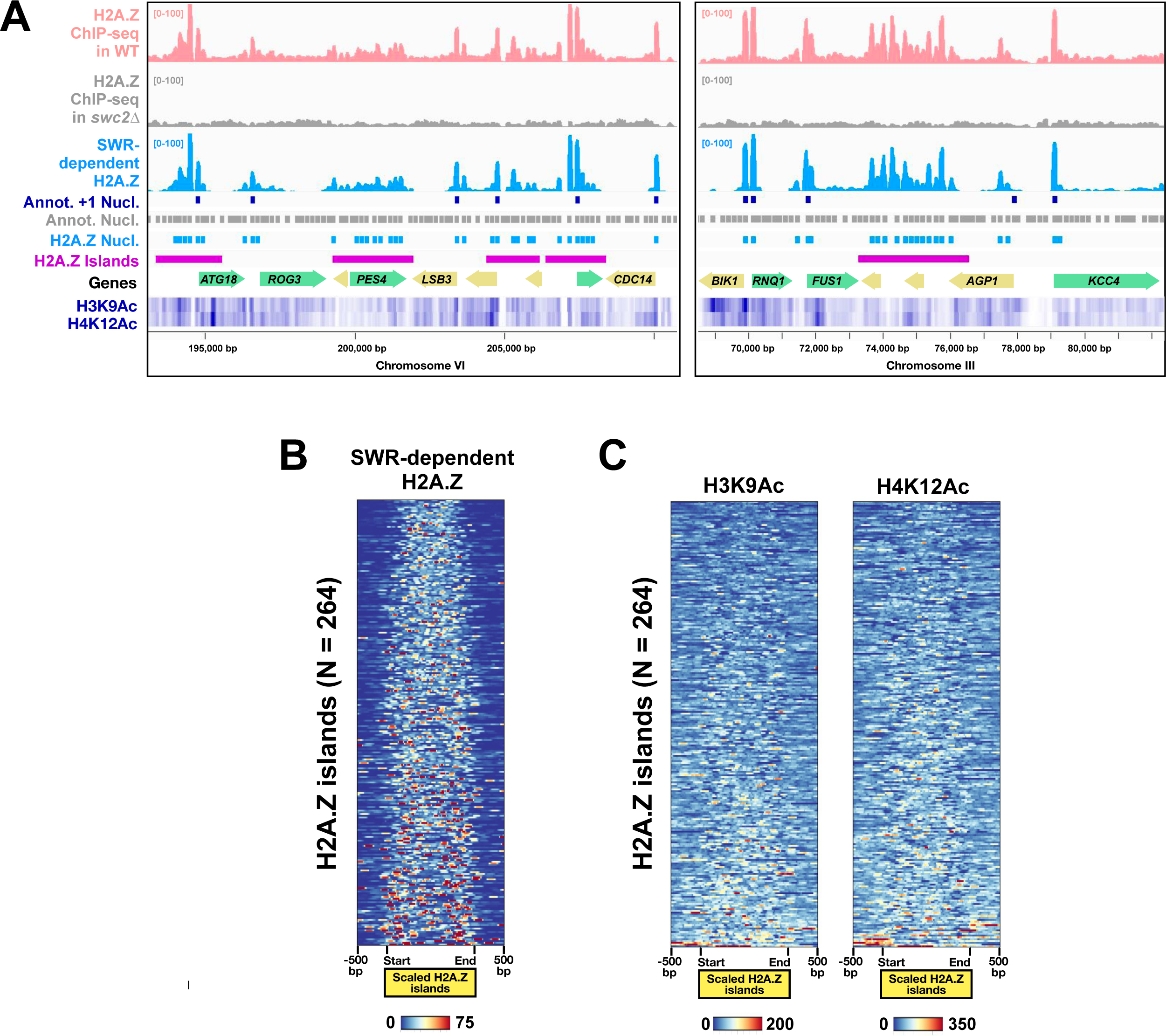
H2A.Z islands. **(A)** Same as 1D except that two other regions with H2A.Z islands were shown. **(B)** A heatmap showing SWR-dependent H2A.Z levels of 264 H2A.Z islands. H2A.Z islands were aligned at their starts and ends, scaled to equal length, and sorted by H2A.Z levels. **(C)** Heatmaps showing H3K9Ac and H4K12Ac levels of the 264 H2A.Z islands.

**Figure S3.**
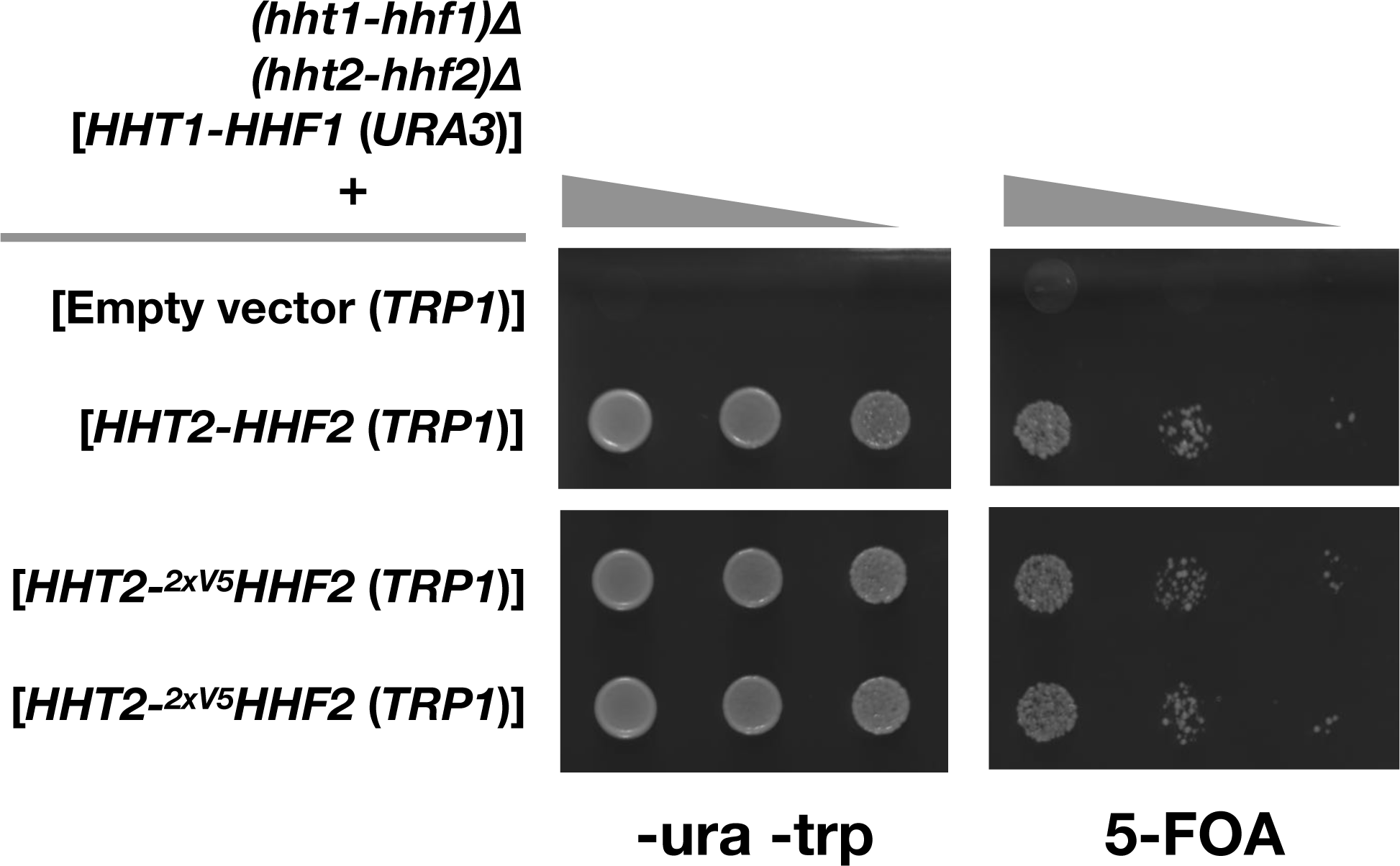
Complementation test showing that the *2xV5-HHF2* gene is functional. The *HHT2-(2xV5-HHF2) TRP1 CEN ARS* plasmid, the untagged control or the empty vector were transformed into a yeast strain that lacked the endogenous H3 and H4 genes but was kept alive by a wild-type *HHT1-HHF1 URA3 CEN ARS* plasmid. Ten-fold serially diluted cells (starting at 1 OD_600_) were spotted onto synthetic complete media lacking uracil and tryptophan (left) or media supplemented with 5-FOA (right).

**Figure S4.**
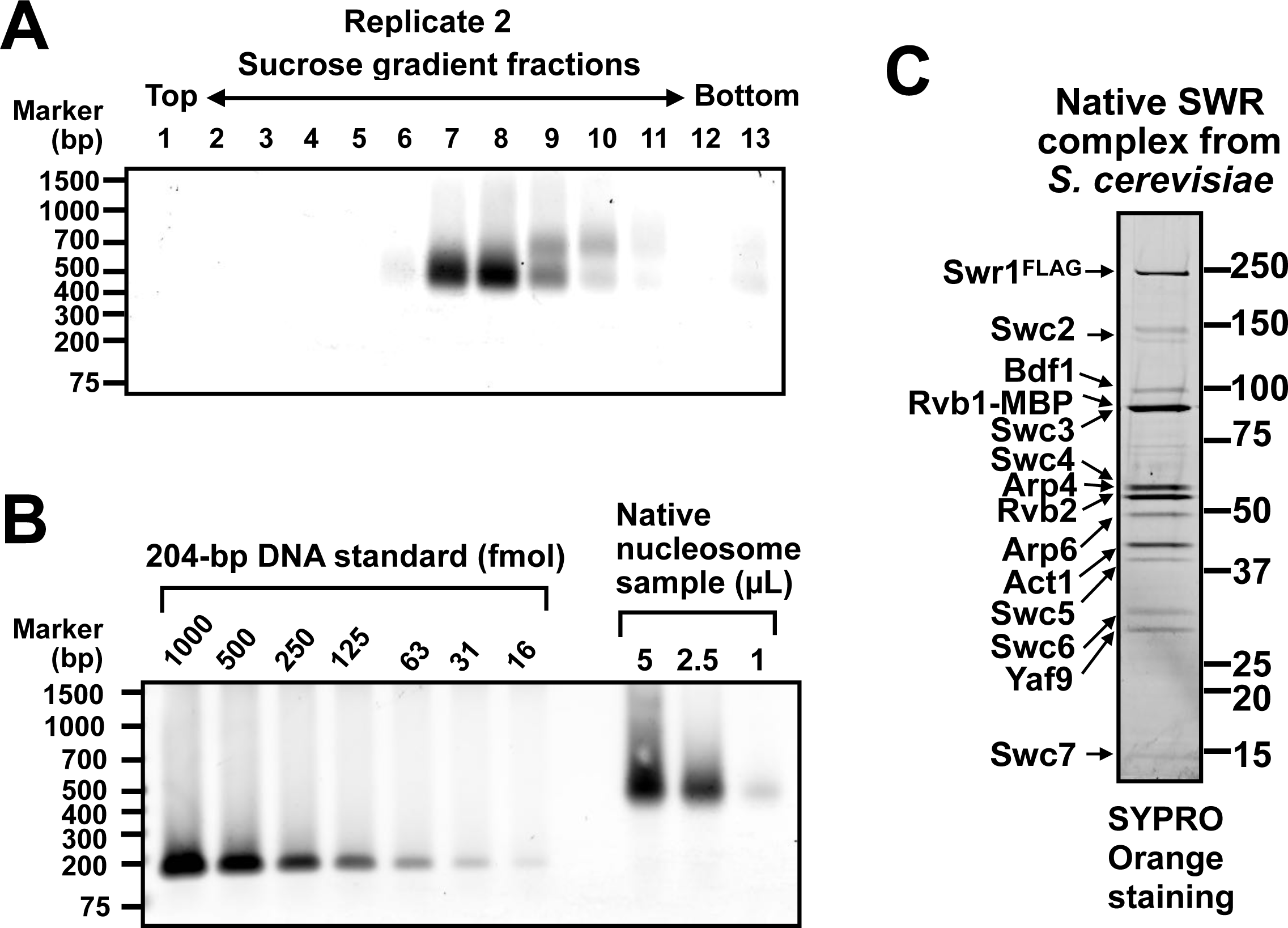
Quality control of the reagents used in the histone exchange reaction. **(A)** Sucrose gradient sedimentation of the anti-V5 affinity purified nucleosomes. This was a replicate of the native nucleosome preparation as shown in Figure 2C. **(B)** Quantification of the native nucleosome preparation. A DNA standard consisting of a 204-bp serially diluted DNA was used to calibrate SYBR gold staining of a native nucleosome samples. **(C)** Native SWR complex purified by the ASAP technique was separated on a 8-16% polyacrylamide gel and analyzed by SYPRO orange staining.

**Figure S5.**
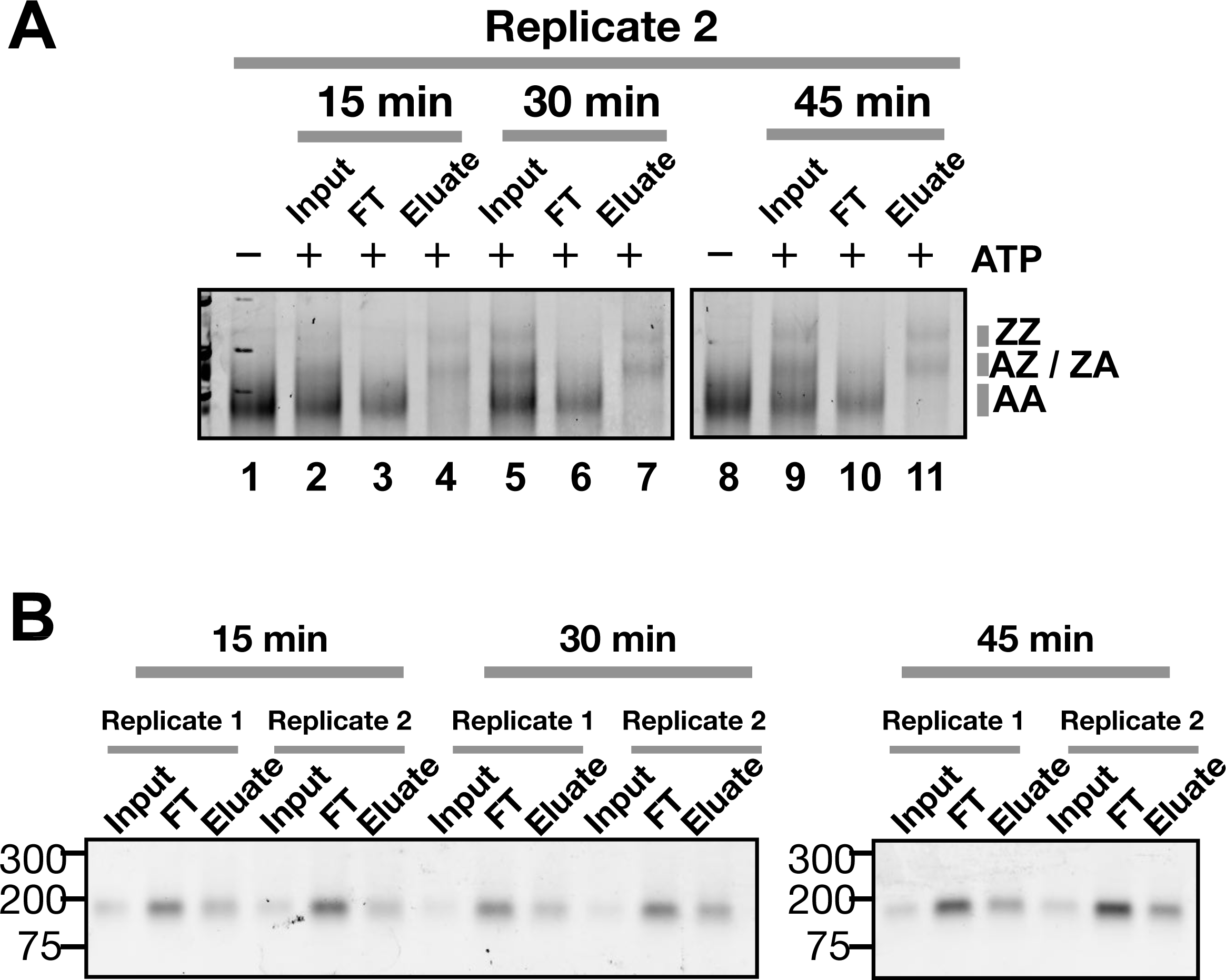
Quality control of the streptavidin pulldown of SWR-remodeled nucleosomes. **(A)** Same as Figure 3C (replicate 1) but a different batch of native nucleosomes were used in replicate 2. **(B)** The nucleosomal DNA from the indicated fractions were extracted and analyzed by a 1.3% agarose / 0.5x TBE gel and SYBR green staining. Sequencing libraries were prepared from the DNA in the FT and eluate fractions.

**Figure S6.**
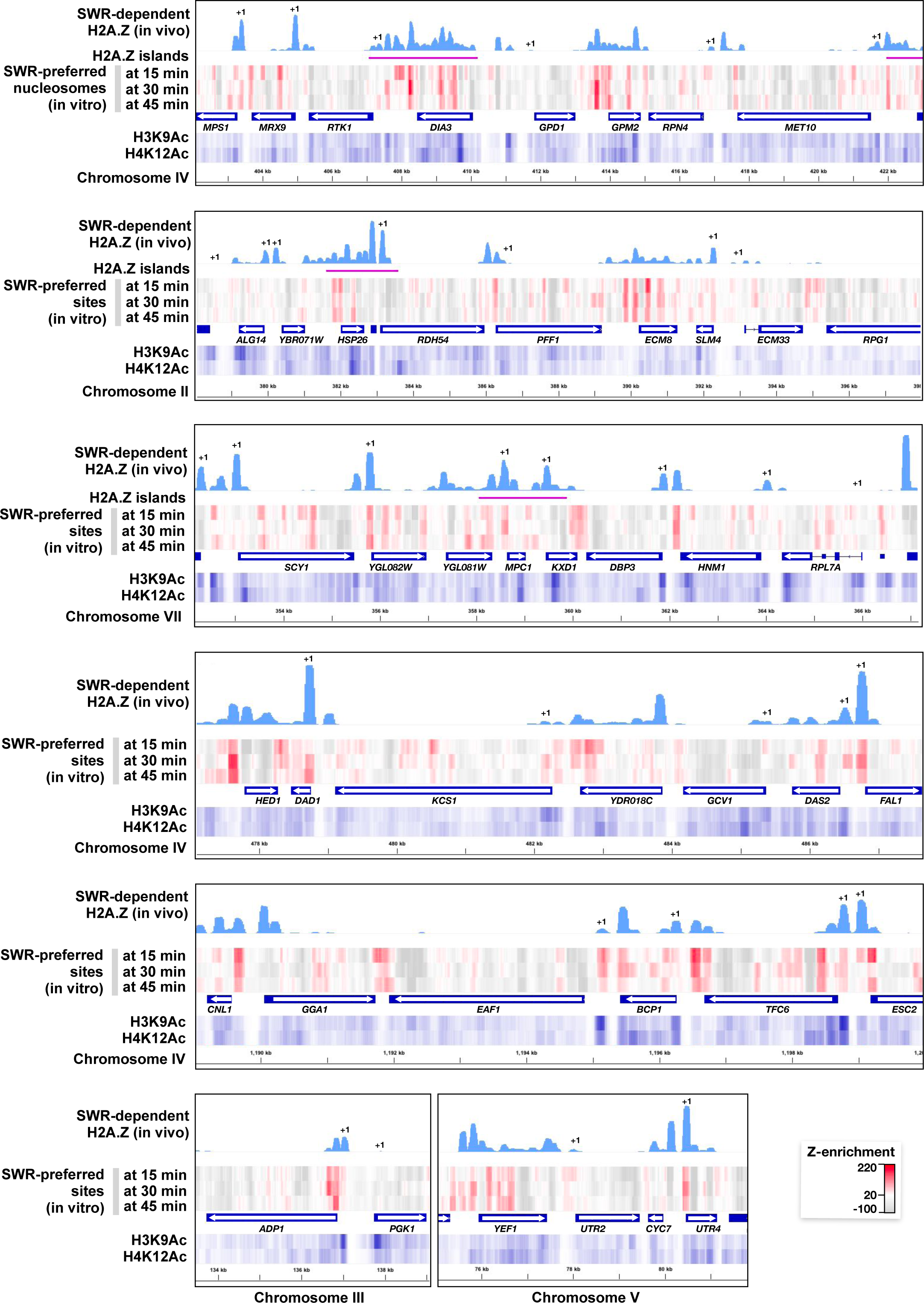
Genomic distribution of SWR-preferred sites. Additional regions plotted in the format of Figure 5A were shown.

**Figure S7.**
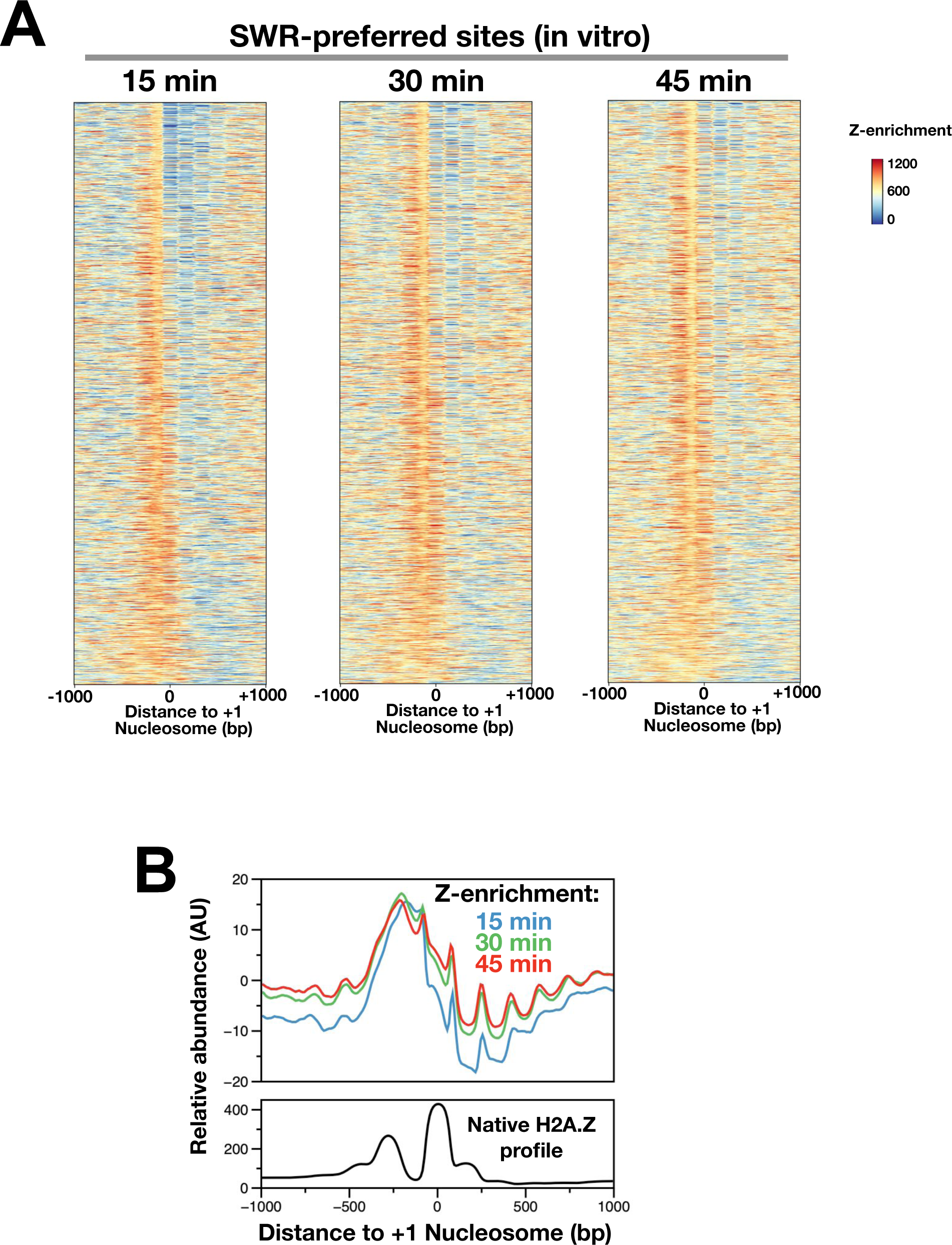
Genomic distribution of SWR-preferred sites in and around +1 nucleosomes. **(A)** Z-enrichment values were centered at +1 nucleosome dyads and sorted using the gene list in Figure 1F left panel. **(C)** Average Z-enrichment values of the heatmaps in A were compared to the native SWR-dependent H2A.Z profile.

**Figure S8.**
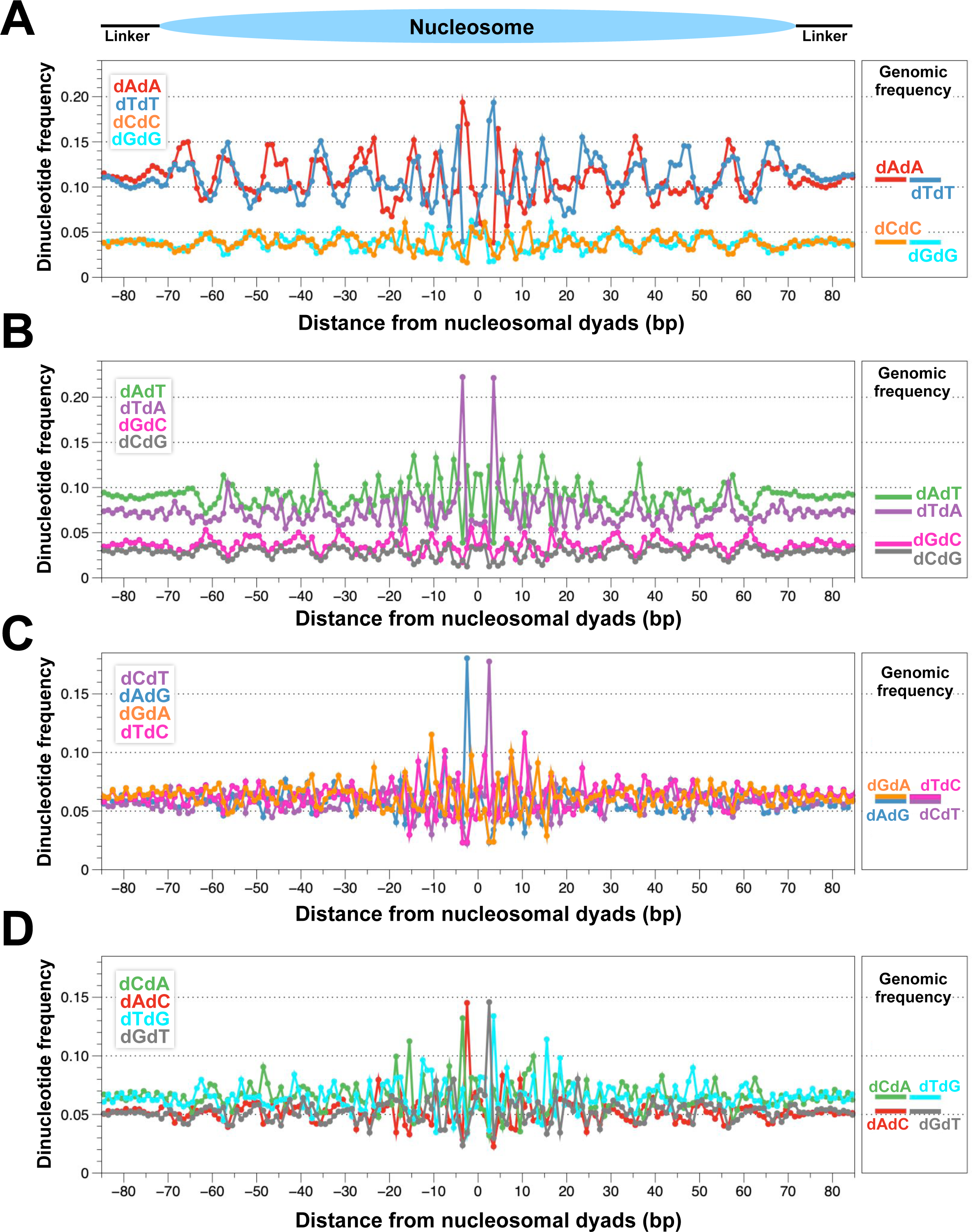
Dinucleotide motif analysis. **(A)** Dinucleotide frequencies of dAdA, dTdT, dCdC, and dGdG were plotted along nucleosome positioning sequences (N = 67,534) centered at nucleosomal dyads plus 20 bp of flanking regions. The box on right shows the frequencies of the dinucleotides across the genome. **(B-D)** same as A, except that the indicated dinucleotide frequencies were shown.

**Figure S9.**
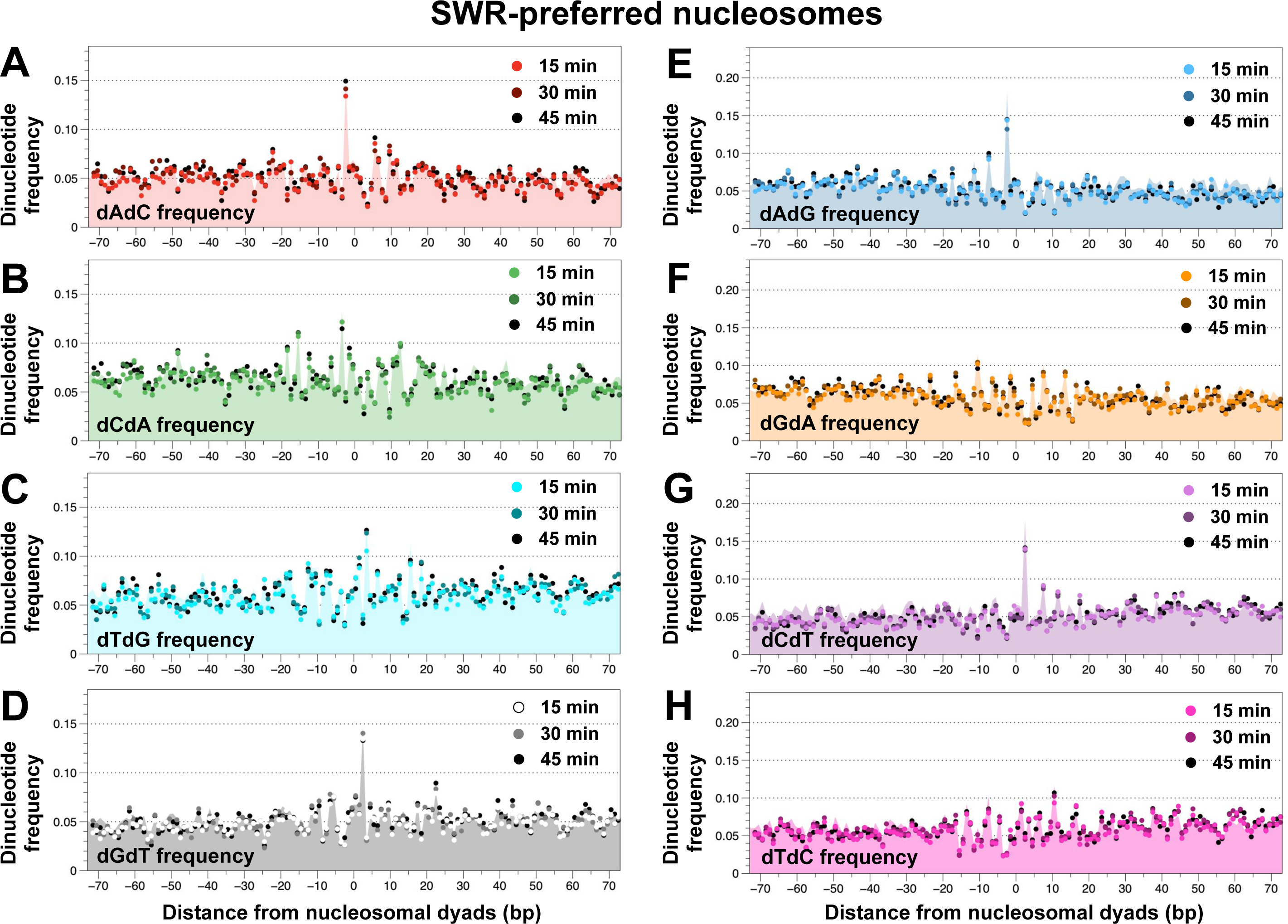
Dinucleotide motif analysis of SWR-preferred nucleosomes. (**A-H**) Same as Figure 6 except that other dinucleotides were shown.

**Figure S10.**
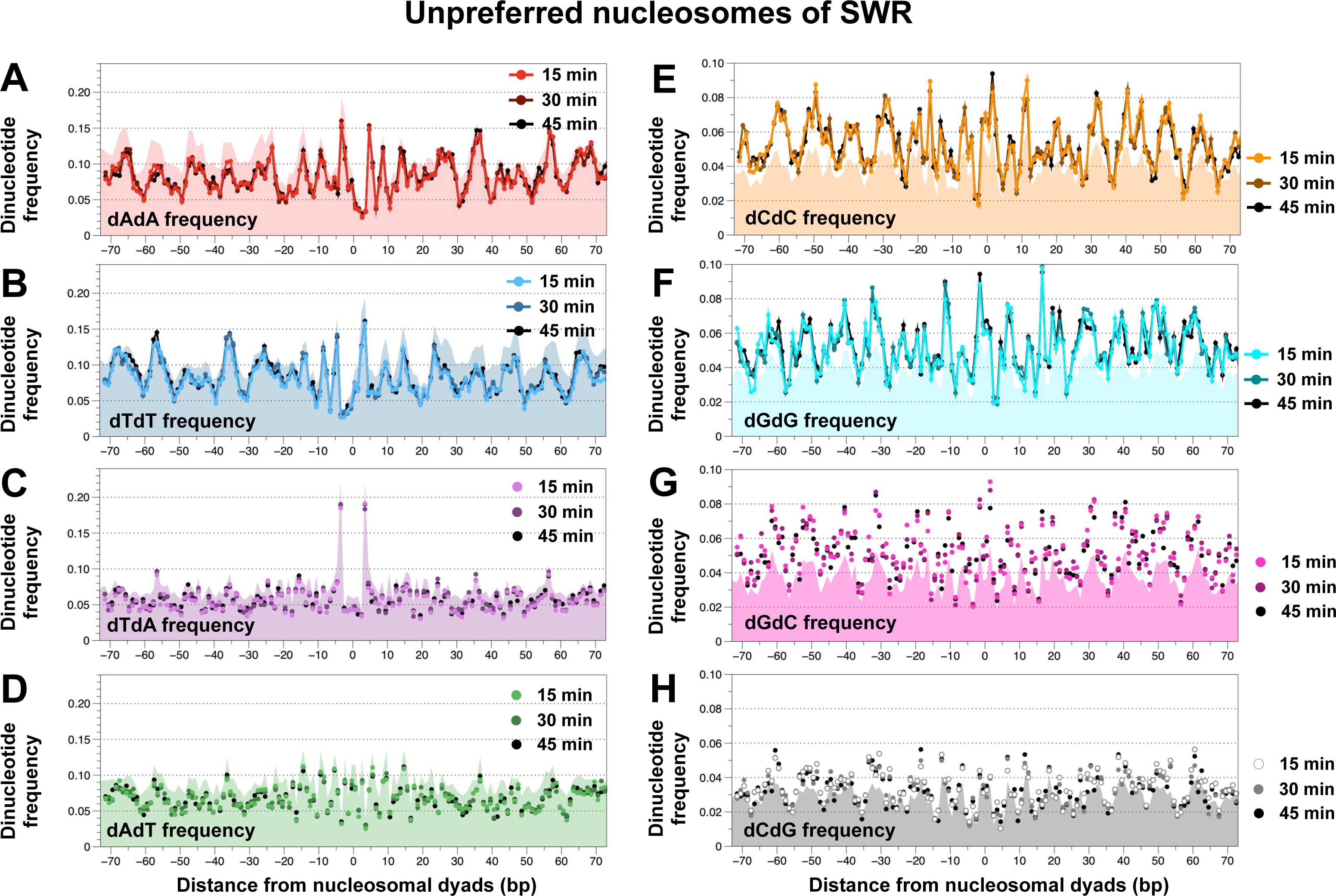
Dinucleotide motif analysis of nucleosomes unpreferred by SWR. (**A-H**) Same as Figure 6 except that the bottom 3% (N = 2,023) of SWR-preferred nucleosomes were used.

**Figure S11.**
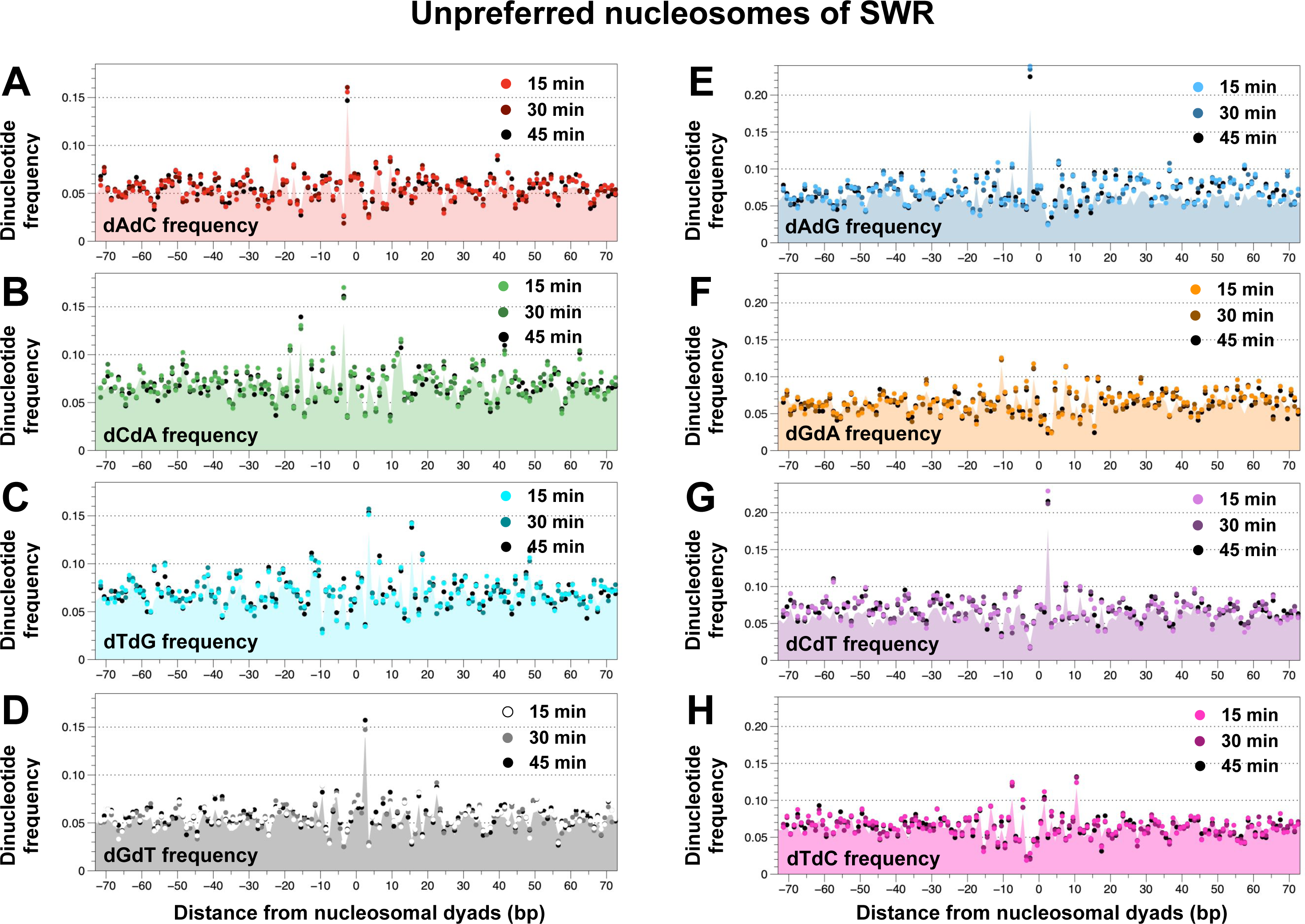
Dinucleotide motif analysis of nucleosomes unpreferred by SWR. (**A-H**) Same as Figure S10 except that the indicate dinucleotide frequencies were plotted.

**Figure S12.**
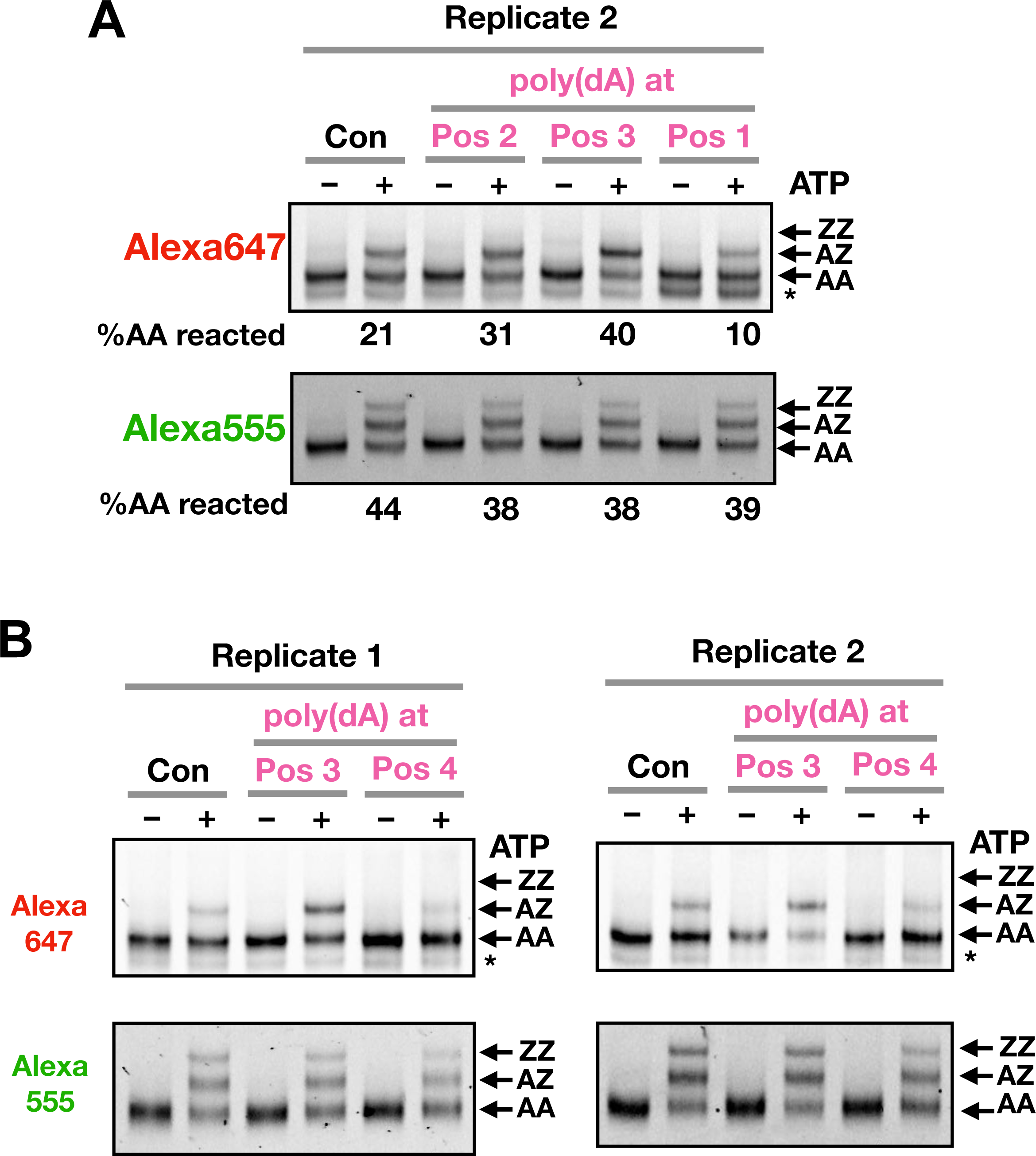
Effects poly(dA) tracks on H2A.Z deposition. **(A)** A replicate of the experiment in Figure 7D. **(B)** Same as A, except that a nucleosomal substrate with a poly(dA) track at Pos 4 was included.

**Figure S13.**
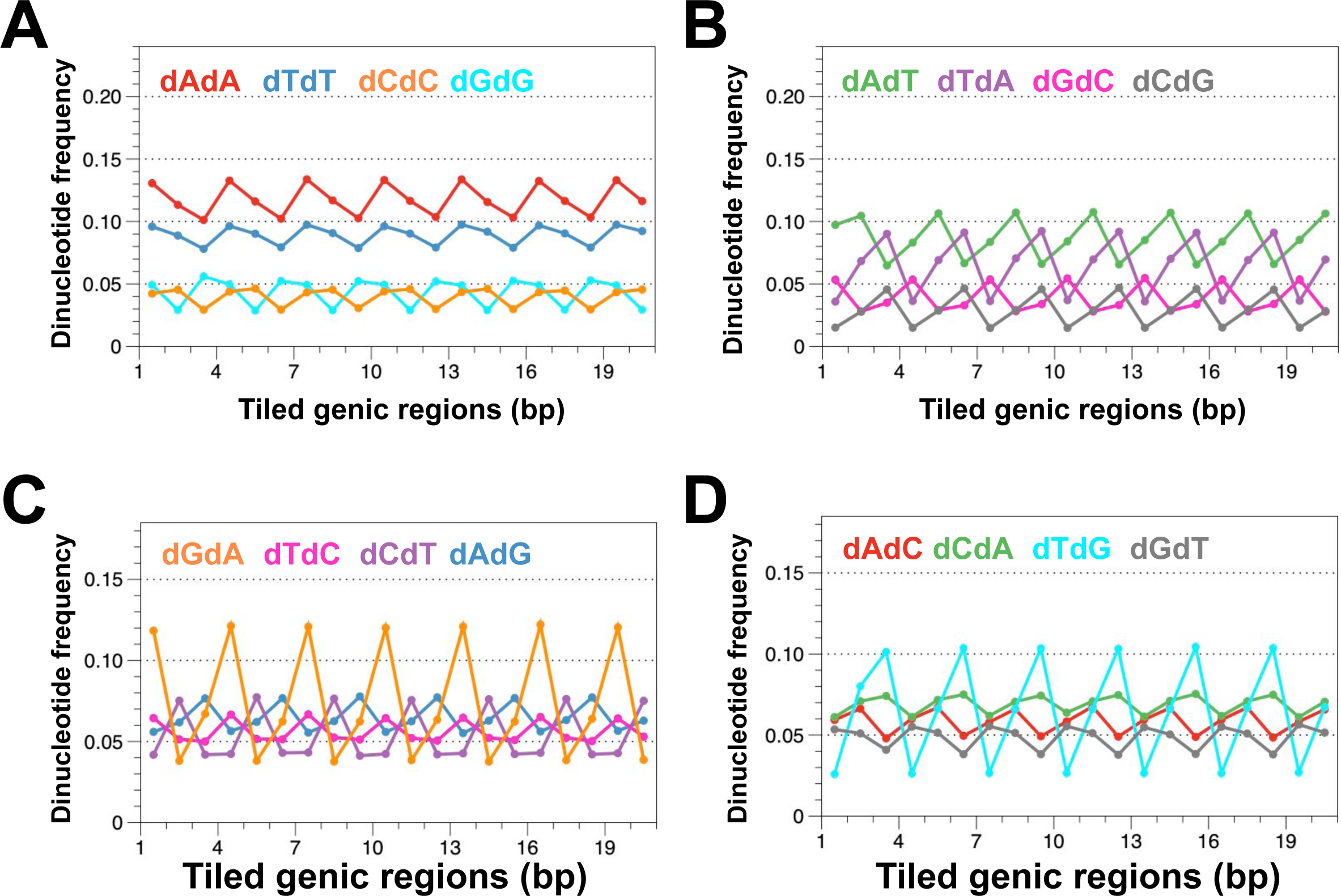
Dinucleotide motif analysis of genic regions. **(A-D)** The coding regions of 6,401 yeast genes were divided into 21-bp tiled fragments in-frame with the genetic codes. The indicated dinucleotide frequencies were averaged over 422,795 tiled regions.

**Figure S14.**
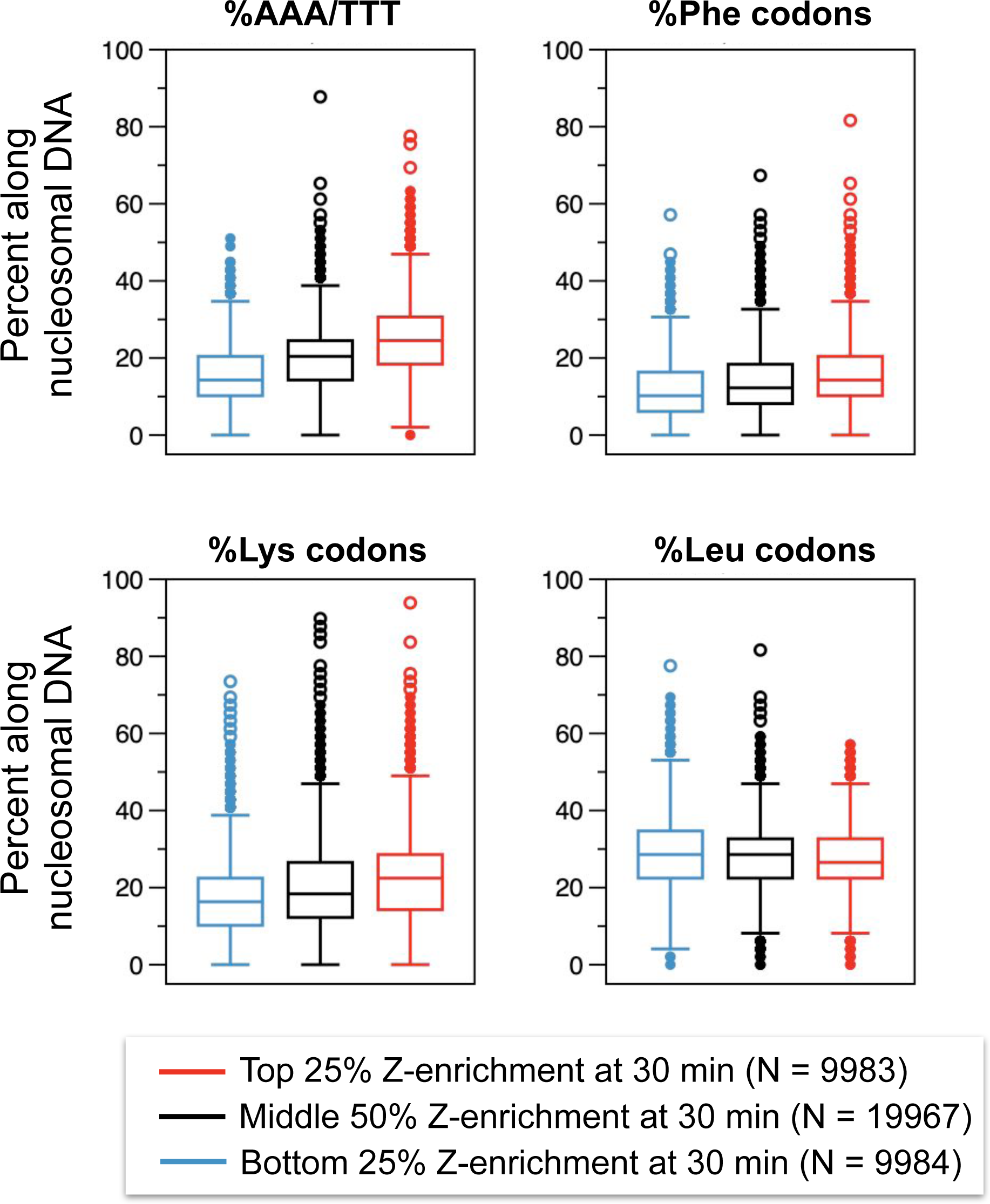
Codon frequencies in SWR-preferred and unpreferred regions.

